# Microevidence for microdosing with psilocybin mushrooms: a double-blind placebo-controlled study of subjective effects, behavior, creativity, perception, cognition, and brain activity

**DOI:** 10.1101/2021.11.30.470657

**Authors:** Federico Cavanna, Stephanie Muller, Laura Alethia de la Fuente, Federico Zamberlan, Matías Palmucci, Lucie Janeckova, Martin Kuchar, Carla Pallavicini, Enzo Tagliazucchi

## Abstract

The use of low sub-hallucinogenic doses of psychedelics (“microdosing”) has gained popularity in recent years. Although anecdotal reports claim multiple benefits associated with this practice, the lack of placebo-controlled studies limits our knowledge of microdosing and its effects. Moreover, research conducted in laboratory settings might fail to capture the motivation of individuals engaged in microdosing protocols. We recruited 34 individuals planning to microdose with psilocybin mushrooms (*Psilocybe cubensis*), one of the materials most frequently used for this purpose. Following a double-blind placebo-controlled design, we investigated the effects of 0.5 g dried mushrooms on subjective experience, behavior, creativity, perception, cognition, and brain activity. The reported acute effects were significantly more intense for the active dose compared to the placebo, which could be explained by unblinding. For the other measurements, we observed either null effects or a trend towards cognitive impairment and, in the case of EEG, towards reduced theta band spectral power. Our findings support the possibility that expectation effects underlie at least some of the anecdotal benefits attributed to microdosing with psilocybin mushrooms.

## Introduction

Over the last decade, the use of relatively small doses of psychedelics has increased in popularity, attracting a significant amount of interest from the general public (Glatter, 2015; Nye, 2017; Sahakian et al., 2018) and the scientific community (Ona, 2020). The Global Drug Survey reported a comparatively high prevalence (28.6%) of this practice, frequently known as “microdosing”, with similar percentages found by other independent studies (Winstock, 2018; Cameron et al., 2020). Microdosing is frequently undertaken to improve mood, cognitive function and mental concentration, as well as to enhance creativity and problem-solving skills (Hutten et al., 2019a; Polito and Stevenson, 2019; Lea et al., 2020). Also, some individuals microdose to self-medicate for cluster headaches, depression and anxiety, among other conditions (Andersson et al., 2017; Hutten et al., 2019b; Lea et al., 2020). Indeed, it has been proposed that microdosing with psychedelics could have therapeutic value for the treatment of these mental health disorders (Kuypers, 2020). The use of low doses of psychedelics constitutes an attractive therapeutic model, since it could circumvent the potential issues associated with altered consciousness and challenging experiences elicited by higher doses (Olson, 2020a).

Owing to its origin as an underground practice, microdosing lacks standards and procedures that are generally accepted and replicated in the community (Kuypers et al., 2019). Different serotonergic psychedelics are used for this purpose, such as lysergic acid diethylamide (LSD), dimethyltryptamine (DMT) ingested with a monoamine oxidase inhibitor (MAOI) —as in the concoction known as ayahuasca— and psilocybin, the active compound of several mushrooms in the *Psilocybe* genus. The most frequently used compounds are LSD and psilocybin, the latter in the form of dried psychoactive mushrooms (Hutten et al., 2019; Polito and Stevenson, 2019; Lea et al., 2020; Szigeti et al., 2021). There is considerable variability in dose and dosing schedules (Kuypers et al., 2019). In the case of psilocybin mushrooms, microdosing is carried out mainly according to four different protocols. Perhaps the most popular schedule was proposed by James Fadiman, consisting of two consecutive days of dosing followed by two days without dosing (Fadiman, 2011). The second approach involves dosing on weekdays, from Monday to Friday, without dosing on Saturdays and Sundays. A third approach is based on dosing two out of every three days (Polito & Stevenson, 2019). Finally, some users dose every day. Dosing periods are highly variable, ranging between one week and several years (Kuypers et al., 2019). In the case of psilocybin mushrooms, microdoses are within the range of 0.1 g to 0.5 g of dried mushroom material (Prochazkova et al., 2018), with 0.1 g considered roughly equivalent to ≈4.6 μg of LSD (Szigeti et al., 2021).

The efficacy of microdosing to enhance mood, creativity and cognition and to reduce anxiety and depression is supported by anecdotal accounts (Fadiman, 2011) and, more recently, by online surveys, observational, and open-label field studies (Johnstad, 2018; Prochazkova et al., 2018; Hutten et al., 2019; Polito and Stevenson, 2019; Anderson et al., 2019a; Anderson et al., 2019b; Fadiman and Korb, 2019; Webb et al., 2019; Cameron et al., 2020; Lea et al., 2020). Unfortunately, these studies lack adequate controls and are based on self-selected samples, rendering them vulnerable to confirmatory bias. It is important to note that expectations (which are generally positive in the context of recent scientific studies) play an important role in the perceived effects of microdosing with psychedelics, both for researchers and participants (Polito and Stevenson, 2019; Olson et al., 2020b; Kaertner et al., 2021; Rootman et al., 2021). When restricted to studies that follow double-blind and placebo-controlled experimental designs, considerably less evidence supports the positive effects of microdosing. Indeed, studies of low LSD doses found effects that are different (or even opposite) to those expected by individuals who microdose (Bershad et al., 2019; Yanakieva et al., 2019; Hutten et al., 2020; Family et al., 2020). Nevertheless, other reports have documented positive and dose-dependent enhancements in mood, emotional cognition and aesthetic perception, as well as significant improvements in emotional state, anxiety, energy and creativity, among other relevant variables (Bershad et al., 2020; Hutten et al., 2020; Szigeti et al., 2021; Van Elk et al., 2021). Importantly, some of these results can be are explained by unblinding, i.e. by subjects correctly distinguishing the placebo from the active condition (Bershad et al., 2020; Van Elk et al., 2021; Szigeti et al., 2021).

There is an important relationship between contextual factors related to the individual (“set”) and the environment (“setting”), and the effects of serotonergic psychedelics (Carhart-Harris et al., 2018). These factors could play an especially important role in studies of microdosing. Both personal motivation and the surrounding environment are relevant variables in the process of producing new and useful ideas (Amabile, 1985; Prabhu et al., 2008). Studies lacking a control condition (i.e. open label studies) usually investigate subjects invested in self-managed microdosing protocols, and therefore have greater ecological validity than randomized, double-blind and placebo-controlled designs. On the other hand, the results of these studies are vulnerable to be distorted by expectation, in particular considering the pro-suggestibility effects of serotonergic psychedelics and the self-reported nature of the measurements (Carhart-Harris et al., 2018; Muthukumaraswamy et al., 2021). While several studies have investigated microdosing in natural settings, except for few exceptions they lacked the controls necessary to avoid confounding by placebo effects and confirmation bias.

In spite of several recent studies addressing the effects of microdosing on mental health, mood, creativity and cognition, physiological and neurobiological levels remain comparatively underinvestigated. Indeed, Kuypers and colleagues highlighted the value of investigating physiological variables, such as heart rhythm and sleep-wake patterns, which could be indicative of negative effects caused by low and repeated doses of 5-HT_2A_ receptor agonists (Kuypers et al. al., 2019). In terms of brain activity, a study using functional magnetic resonance imaging (fMRI) showed that a very low dose of LSD sufficed to alter the functional connectivity between the amygdala and several cortical regions; moreover, some of these changes were correlated with self-reported assessments of positive mood (Bershad et al., 2020). Besides fMRI, other non-invasive techniques show potential to map the effects of serotonergic psychedelics in large-scale brain activity; in particular, electroencephalography (EEG) is capable of robustly identifying the acute effects of different psychedelics by detecting broadband increases in signal entropy and band-specific changes in spectral power (Riba et al., 2002; Muthukumaraswamy et al., 2013; Kometer et al., 2013; Schenberg et al., 2015; Schartner et al., 2017; Valle et al., 2016; Carhart-Harris et al., 2016; Timmermann et al., 2019; Pallavicini et al., 2021; Tagliazucchi et al., 2021). Despite this promising previous research, to date only one study applied EEG to investigate the effects of low doses of LSD, finding dose-dependent reductions in broadband oscillatory power during resting state with eyes open and closed, as well as modulation of event-related potentials (ERPs) in a visual oddball paradigm (Murray et al., 2021).

The present study investigated the effects of low doses of *Psylocybe cubensis* on behavior, creativity, perception, cognition and the underlying brain activity, with emphasis on controlling for expectation without introducing an artificial context for microdosing. We recruited individuals who were planning to start a microdosing protocol with their own mushroom material and who willingly adapted their schedule and dose to meet the standardized conditions of our research protocol. The experimental condition (gel capsule with either 0.5 g dried *Psylocybe cubensis* or the same weight of inactive placebo) was unknown to both participants and experimenters, and was only revealed after data collection and analysis was concluded, in accordance with a randomized double-blind placebo-controlled experimental design.

## Materials and methods

### Recruitment and participants

In total, 34 participants (11 females; 31.26 ± 4.41 years; 74 ± 17 kg [mean±STD]) were recruited by word-of-mouth, social media advertising, and visits to psilocybin mushroom and microdosing workshops between December 2020 and August 2021. Participants reported 11 ± 14.9 previous experiences with serotonergic psychedelics, of which 1.5 ± 2.3 were considered “challenging”; only 6 of them reported significant previous experience with microdosing. All participants were fluent in Spanish, had normal or corrected-to-normal vision, and successfully completed all instances of the experiment. Interested individuals were asked to get in contact with the researchers by email, which was followed by a phone conversation. During this first contact, candidate participants were briefed on the details and objectives of the experiment, as well as on the inclusion and exclusion criteria. They were given a copy of the informed consent form together with a detailed written explanation of the experiment and its objectives. After signing the form, all subjects underwent a psychological interview to screen for the exclusion criteria that are detailed later in this section. Afterwards, participants and researchers agreed on the start of the microdosing procedure, which followed the schedule presented in the next session. A multi-sensor commercial sleep and activity-tracker wristband (Fitbit Charge 4) was sent to all participants at the beginning of each week of the study. During the experiment, each individual was assessed on four different days by the research team (two days for the active dose and two for the placebo, see “Design” subsection), which consisted of four members, and always included a clinical psychologist (FC).

This study was conducted in accordance with the Helsinki declaration and approved by the Committee for Research Ethics at the Universidad Abierta Interamericana (Buenos Aires, Argentina), protocol number 0-1054. The experiments entailed no deception and participants were fully informed about the purpose of the study. After the study ended, the order of the conditions was unblinded to the subjects. The participants did not receive financial compensation for their participation.

### Participant selection criteria

Individuals who at the time of contact were planning to start a microdosing protocol with their own *Psilocybe cubensis* material were contacted and informed about the experimental design, and were given information about the proposed dosing scheduling, with a detailed explanation of the need for a blinded and randomized control condition with an inactive placebo. To be included as participants of this experiment, subjects were asked to comply with the schedule of the experiment (full details in a following subsection), to abstain from consuming psychoactive drugs (including alcohol and caffeine) during the study weeks, and to avoid eating three hours before consuming each microdose. A non-diagnostic psychiatric interview was conducted according to the guidelines by Johnson et al. (2008). Subjects who fulfilled DSM-5 criteria for the following disorders were excluded from the experiment: schizophrenia or other psychotic disorders, and type 1 or 2 bipolar disorder (both also in first and second degree relatives), substance abuse or dependence over the last 5 years (excluding nicotine), depressive disorders, recurrent depressive episodes, obsessive-compulsive disorder, generalized anxiety disorder, dysthymia, panic disorder, bulimia or anorexia, as well as subjects with history of neurological disorders. Subjects under psychiatric medication of any kind were also excluded. All participants provided their written informed consent to participate in the study.

### Blinding procedure and experimental design

The design, outlined in Figure 1, was structured as follows. For each participant, the experiment was divided into two consecutive weeks, one corresponding to the active dose (0.5 of finely ground and homogenized dried mushroom material in a gel capsule) and the other to the placebo (same weight of a dried edible mushroom). A dose of 0.5 g is representative of the upper range of values used in microdosing (Prochazkova et al., 2018; Polito and Stevenson, 2019; Van Elk et al., 2021; Szigeti et al., 2021). The order of these two conditions was randomized by a third party and the identity of the capsules was unknown both to the participant and the researchers. This procedure is similar to the one implemented in a recent publication from the Center for Psychedelic Research, Imperial College, London (https://selfblinding-microdose.org/; Szigeti et al., 2021), except that blinding was not performed by the participants themselves, thus avoiding the possibility of its deliberate breaking (Van Elk et al., 2021). The complete procedure consisted of the following steps:

1. Two gel capsules were filled with 0.5 g of dried and finely ground and homogenized *Psilocybe cubensis*, yielding two active doses of 0.5 g each. These capsules were stored in an airtight plastic bag within a paper envelope. For each independent source of mushrooms, samples of 150 mg were taken and preserved for chemical analysis.
2. This process was repeated using the same number of capsules, but each filled with 0.5 g of dried and ground edible mushrooms (e.g. *Suillus granulatus*). These capsules were stored in an airtight plastic bag within paper envelopes identical to those used in the previous step.
3. The blinded conditions could be identified by a folded paper with a code that was introduced to both envelopes.
4. Subjects took the envelopes and randomly selected one of them at the beginning of the experiment, leaving the other for the second week.
5. After the data was collected, scored and analyzed, subjects reported to the experimenters the code corresponding to each of their envelopes. Next, the third party in charge of the blinding used these codes to determine which envelope corresponded to each condition, sharing this information with the researchers to perform the final statistical analyses. In turn, the order of the conditions was also revealed to the participants by the researchers.

**Figure 1.**
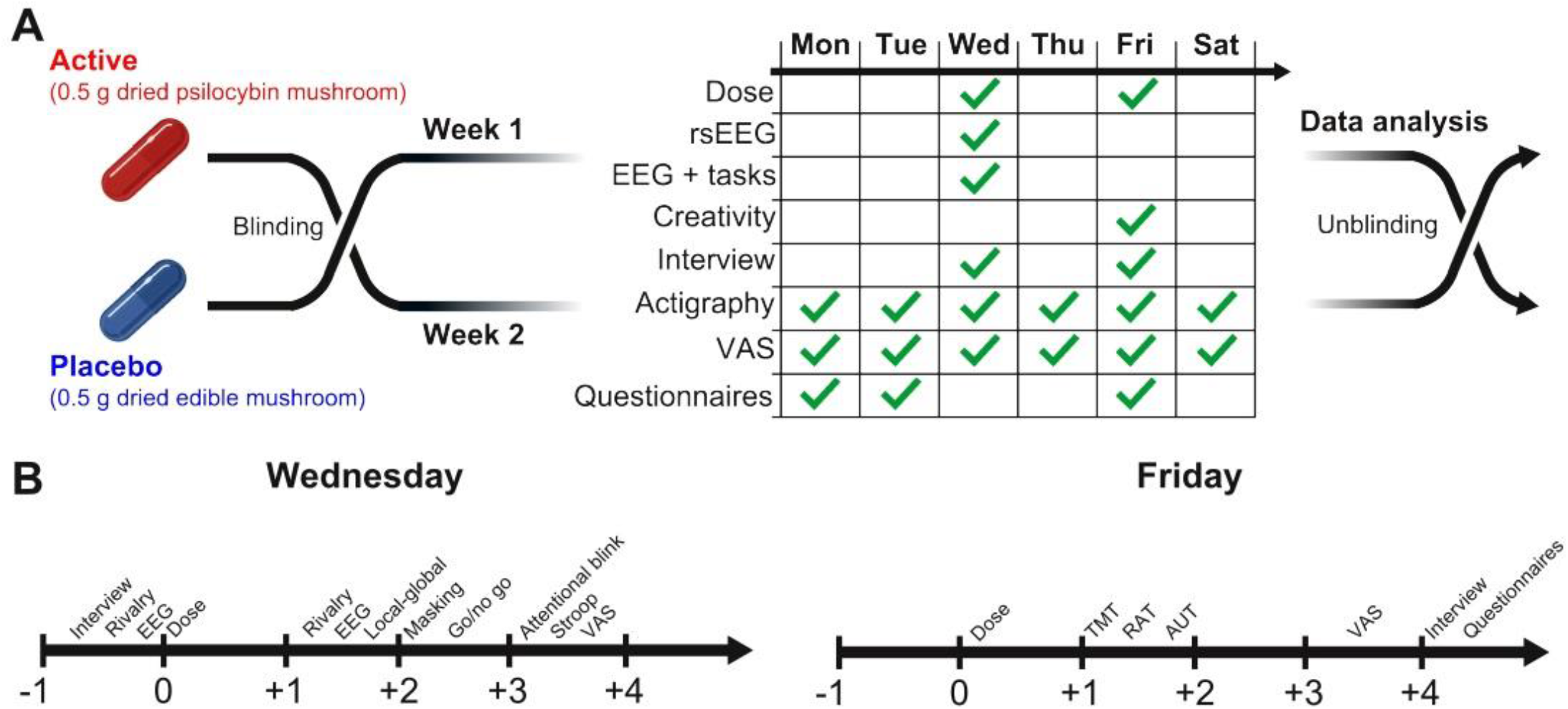
Experimental design. A) Double-blind placebo-controlled design implemented to mask the active condition. Neither the subjects nor the investigators knew the content of the capsules until the last steps of the data analysis stage. Each condition (active dose or placebo) corresponded to one different week of the experiment (separated by one week), with different measurements conducted depending on the day, starting ≈1.5 h after consuming the capsule (panel A, right). B) Wednesday (first dosing day) and Friday (second dosing day) comprised two different sets of measurements, indicated in the corresponding timelines (“0” indicates the time when subjects consumed their capsules).

Baseline measurements of psychological traits were conducted on the first day of the week for each condition. Also, participants received and started wearing the Fitbit Charge 4 wristband to track their daily levels of physical activity. On all the days of the protocol, subjects also completed a self-reported scale to assess the subjective effects perceived during the day, accessed through a link that they received via Telegram.

Capsules (active or placebo, depending on the week) were consumed on Wednesday (first dosing day) and Friday (second dosing day). On the same days, the participants visited the experimental premises and performed a series of measurements that are detailed in one of the following sections. Measurements started ≈1.5 h after consuming the capsule (Passie et al., 2002).

### Experimental setting

All experiments were conducted in a comfortable house which hosted only the researchers and the participant, with separate rooms fitted for the needs of each task. Environmental sounds were kept to a minimum and participants were provided with noise-cancelling headphones in case they were necessary. During EEG recordings, the main power line was interrupted in order to avoid artifacts due electrical currents in the proximity of the electrodes. To avoid the possibility of subjects driving under the effects of psilocybin, all subjects were taken by car to the experimental premises and then back to their points of departure

Participants completed a series of self-reported scales aimed to assess psychological traits and expectations two days before each condition. Afterwards, they performed different tasks and activities on Wednesdays and Fridays, and completed a battery of psychometric scales on Fridays. Table 1 summarizes all the tasks and measurements included in this study. A detailed description of each task and measurement is in the following subsections.

**Table 1.** Questionnaires, tasks and measurements collected for each condition, including the domain they target, acronyms, and when they were obtained during the experiment (baseline, first dosing day, second dosing day, daily).

| Domain | Instrument / Task | Acronym | Mon.<br>(baseline) | Wed.<br>(1<sup>st</sup> dose) | Fri.<br>(2<sup>nd</sup> dose) | Daily |
| --- | --- | --- | --- | --- | --- | --- |
| Acute effects | Visual Analog Scale | VAS |  | X | X | X |
| Personality | Big Five Inventory | BFI | X |  |  |  |
| Trait anxiety | State Trait Anxiety Inventory | STAI-T | X |  |  |  |
| Suggestibility | Short Suggestibility Scale | SSS | X |  |  |  |
| State anxiety | State Trait Anxiety Inventory | STAI-S |  |  | X |  |
| State affect | Positive and Negative Affect Scale | PANAS |  |  | X |  |
| Stress | Perceived Stress Scale | PSS |  |  | X |  |
| Absorption | Tellegen Absorption Scale | TAS |  |  | X |  |
| Well-being | Well-being Scale | BIEPS |  |  | X |  |
| Mind wandering | Mind Wandering Questionnaire | MWQ |  |  | X |  |
| Flow | Flow State Scale | FSS |  |  | X |  |
| Creativity | Creative Personality Scale | CPS |  |  | X |  |
| Empathy | Cognitive-Affective Empathy Test | TECA |  |  | X |  |
| Cognitive flexibility | Cognitive Flexibility Scale | CFS |  |  | X |  |
| Convergent thinking | Remote Associates Test | RAT |  |  | X |  |
| Divergent thinking | Alternative Uses Task | AUT |  |  | X |  |
| Divergent thinking | Wallach-Kogan Test | WK |  |  | X |  |
| Perception | Binocular Rivalry | BR |  | X |  |  |
| Conscious perception | Backward Masking | BM |  | X |  |  |
| Attention, coordination | Trail Making Test | TMT |  |  | X |  |
| Inhibition | Go / No Go | GNG |  | X |  |  |
| Attention | Attentional Blink | AB |  | X |  |  |
| Inhibition | Stroop Test | ST |  | X |  |  |
| Brain activity | EEG with Eyes Open | EO |  | X |  |  |
| Brain activity | EEG with Eyes Closed | EC |  | X |  |  |
| Attention | Local-Global + EEG | LG |  | X |  |  |
| Physical activity | Fitbit Charge 4 wristband | ACT | X | X | X | X |

### Acute effects

Visual Analog Scale (VAS). The items were translated and adapted from Carhart-Harris et al. (2016) and presented in the form of VAS to determine the intensity of the effects experienced by subjects. The items rated in the VAS were the following: “My imagination was extremely vivid”, “The experience had a dreamlike quality”, “Sounds influenced things I saw”, “My sense of space and size was distorted”, “I felt unusual bodily sensations”, “My thoughts wandered freely”, “My perception of time was distorted”, “I saw geometric patterns”, “Edges appeared warped”, “My thinking was muddled”, “I saw movement in things that weren’t really moving”, “I experienced a sense of merging with my surroundings”, “Things looked strange”, “I felt like I was floating”, “The experience had a supernatural quality”, “I experienced a disintegration of my self or ego”, “I felt a profound inner peace”, “The experience had a spiritual or mystical quality”, “I felt afraid”, “I feared losing control of my mind”, “I felt suspicious and paranoid”).

### Self-reported scales and questionnaires

Big Five Inventory (BFI). An inventory assessing five dimensions of personality: neuroticism, extraversion, openness to experience, agreeableness, and conscientiousness (Benet-Martinez and John, 1998). The BFI questionnaire consists of 44 items based on a 5-point Likert scale.

State-Trait Anxiety Inventory (STAI-T / STAI-S). Commonly used scales which measure state anxiety (situational anxiety of a temporary nature) and trait anxiety (stable trait linked to individual characteristics) (Spielberger et al., 1983). The instrument comprises 40 items and is based on a 4-point Likert scale.

Short Suggestibility Scale (SSS). An inventory that assesses suggestibility, created by Kotov et al. (2004). The questionnaire consists of 21 items based on a 5-point Likert scale.

Positive and Negative Affect Schedule (PANAS). A psychometric scale that has been widely used to measure dimensions of affect, both positive and negative (Watson et al., 1988). The instrument consists of 20 affirmations based on a 5-point Likert scale.

Perceived Stress Scale (PSS). An instrument to assess how different situations affect the feelings and perceived stress of the respondents, consisting of 10 affirmations based on a 5-point Likert scale (Cohen et al., 1994).

Tellegen Absorption Scale (TAS). A 34-item scale developed to measure the capacity of an individual to become absorbed in the performance of a task (Tellegen and Atkinson, 1974).

Psychological Well-being Scale (BIEPS). A scale used to measure eudemonic well-being in adults (including dimensions of acceptance, perception of control, social ties, and autonomy and projects) (Casullo & Brenlla, 2002). It consists of 13 questions based on a 3-point Likert scale.

Mind Wandering Scale (MWQ). *A* 5-question instrument developed by Mrazek et al. (2013) to measure mind-wandering trait levels. It is a 6-point Likert-type scale.

Flow State Scale (FSS). Developed by Jackson & Marsh (1996), the FSS is a 36-item scale, rated on a 5-point Likert-type scale, that measures nine different dimensions of the flow state.

Creative Personality Scale (CPS). The scale is aimed to assess creative behavior and creative personality traits (Gough et al., 1979). The instrument consists of 21 affirmations based on a 4-point scale.

Cognitive-Affective Empathy Test (TECA). Developed by López-Pérez et al. (2008), TECA is a test that aims to measure both components that empathy is comprised of. The test has 33 items in a 5-point Likert scale.

Cognitive Flexibility Scale (CFS). A scale used to measure three factors of cognitive flexibility (Martin & Rubin, 1995). It has 12 items in a 6-point Likert-type scale.

### Creativity tests

Remotes Associates Test (RAT). A test was developed by Mednick & Mednick (1959), widely used to measure creative convergent thinking. The subject is given three words that appear to be unrelated and must think of a fourth word that is related to the previous ones.

Alternative Uses Task (AUT). The test assesses divergent thinking and creativity. Subjects were asked to think of and write of as many uses as possible for a simple item during two minutes (Guilford, 1967).

Wallach-Kogan Test (WK). The test assesses divergent thinking and creativity. Subjects were asked to come up with as many items as possible within a certain general group, without time constrains (Wallach and Kogan, 1965).

### Tasks to measure perception and cognition

Binocular rivalry (BR). A pair of superimposed circles with opposite 45° gratings were presented within 5.4° of the fixation cross. When viewed through red and green filter glasses, each of the gratings was presented to one of the participant eyes. Subjects were instructed to maintain fixation and to report changes of the dominant stimulus by pressing a key, lasting for a total of 10 minutes. Previous reports have shown that the alternation rate between dominant stimuli is modulated by psilocybin (Carter et al., 2007; Carter et al., 2010). This task is outlined in the first row of Figure 5.

Backward masking (BM). Task used to study conscious visual perception, adapted from Del Cul et al. (2007). A digit between 0 and 9 (known as “target”) was briefly flashed (17 ms) at the left/right of the former location of the fixation cross (the location is randomly chosen) using different levels of contrast with the background. After a variable delay, known as stimulus onset asynchrony (SOA), a mask consisting of four letters appeared surrounding the former location of the target. Subjects were asked to indicate whether they perceived the target (subjective visibility report) and then to compare its magnitude relative to number 5 (objective visibility report). First, subject-specific initial contrast values were determined using a staircase procedure, in which the contrast was decreased (i.e. the task was made harder) every time the subjective visibility report was correct, and vice versa. The final contrast was computed as the mean over the 18 last reversals. Afterwards, the subject performs the task with the contrast value found in the staircase procedure with SOA, randomly using the values 16, 32, 478, 64 and 80 ms for the SOA. The performance is given by the objective and subjective visibility accuracy vs. SOA. This task is outlined in the first row of Figure 5.

Trail Making Test (TMT). A widely used test to assess functions such as attention, flexibility, speed and visuomotor integration. Performance is evaluated using two different tracking conditions: Part A involves connecting numbers from 1 to 25 in ascending sequence, while Part B involves connecting numbers and letters in alternating and ascending order (Reitan & Wolfson, 1993). This task is outlined in the first row of Figure 6.

Go / No Go (GNG). This task is designed to measure response inhibition. Subjects were asked to respond as quickly as possible to a ‘go’ stimulus (string “SSSTSSS”) while avoiding to respond to a ‘no go’ stimulus (string “SSSHSSS”). Before presentation, both strings were briefly masked with numerals (i.e. ‘#######’) during 50 ms. The performance was assessed in terms of the response time (RT) and the normalized accuracy (number of trials with correct responses divided by 200). This task is outlined in the first row of Figure 6.

Attentional Blink (AB). Used to measure the refractory period that follows the successful detection of a target in an attention demanding task (Shapiro et al., 1997). On each trial, a fixation cross was presented in the center of a laptop screen for a duration randomly selected between 1000 and 1500 ms. Each fixation cross was followed by a stream of serial visual stimuli consisting of 10–21 black letters, each presented for 100 ms and randomly selected from the alphabet with the following two restrictions: 1) pairs of successive letters could not be the same and 2) letters I, O, Q, S, X and Z were excluded. Two of the letters of the sequence were replaced by red digits between 2 and 9, which corresponded to targets T1 and T2. Targets were separated by lags ranging for 1 position in the sequence (100 ms) to 7 positions (700 ms), with T2 being presented 2 to 4 positions before the end of the stream. At the end of each trial, participants were instructed to detect digits T1 and T2, and their performance was determined by the rate of correct detections of each target. The task consisted of 140 trials, with the relevant parameters randomized across trials. This task is outlined in the first row of Figure 6.

Stroop Test (ST). A neuropsychological test used to assess the ability to inhibit cognitive interference (Stroop, 1935). At each trial, participants were shown a word corresponding to a color (“blue”, “yellow”, “red”, “green”), with a color congruent (e.g. “yellow” shown in yellow) or incongruent (e.g. “yellow” shown in blue) with the word. At the center of the screen, the fixation point was surrounded by one word for each possible color (in randomized positions) and participants were prompted to select the stimulus color using the keyboard arrow keys. The experiment consisted of 50 trials, with the same proportion of congruent and incongruent word-color pairs. Performance in this task was assessed by the normalized accuracy and the RT. This task is outlined in the first row of Figure 6.

### Resting state EEG recording, preprocessing and analysis

EEG was recorded with a 24-channel research-grade mobile system (mBrainTrain LLC, Belgrade, Serbia; http://www.mbraintrain.com/) attached to an elastic electrode cap (EASYCAP GmbH, Inning, Germany;www.easycap.de). Twenty-four Ag/AgCl electrodes were positioned at standard 10–20 locations (Fp1, Fp2, Fz, F7, F8, FC1, FC2, Cz, C3, C4, T7, T8, CPz, CP1, CP2, CP5, CP6, TP9, TP10, Pz, P3, P4, O1, and O2). Reference and ground electrodes were placed at FCz and AFz sites. The wireless EEG DC amplifier (weight = 60 g; size = 82 × 51 × 12 mm; resolution = 24 bit; sampling rate = 500 Hz, 0–250 Hz pass-band) was attached to the back of the electrode cap (between electrodes O1 and O2) and sent digitized EEG data via Bluetooth to a Notebook held by a experimenter sitting behind the participant. EEG activity was acquired with eyes open and closed (five minutes each) and during different tasks (Local-Global, binocular rivalry, Go / No Go, attentional blink, Stroop, backward masking). Here we reported only the results for resting state and the Local-Global auditory stimulation paradigm (see one of the following subsections for a detailed description).

EEG data was preprocessed using EEGLAB (https://sccn.ucsd.edu/eeglab/index.php) (Delorme and Makeig, 2004). First, time series were bandpass-filtered (1 – 90 Hz) and notch-filtered (47.5 – 52.5 Hz). Channels with artifacts were detected using EEGLAB automated methods, with rejection criteria based on kurtosis (threshold = 5), probability (threshold = 5) and the rejection of channels with ±2.5 standard deviations from the mean in any parameter (rejected channels for eyes open, psilocybin: 2±1; eyes closed, psilocybin: 1.5±0.9; eyes open, placebo: 2±1; eyes closed, placebo: 1.5±0.9 [mean±STD]). Channels were manually inspected before rejection and then interpolated using data from the surrounding channels. Next, time series were divided into 2 s epochs, and the first 5 volumes were discarded. Rejected epochs were flagged automatically and visually inspected before rejection. Infomax independent component analysis (ICA) was applied to data from each individual participant with the purpose of identifying and removing components related to eye movements, blinking, heartbeat, and muscle artifacts. This semi-automatic procedure was based on the frequency content and scalp topography of each component (Brunner et al., 2013), and was corroborated by visual inspection (removed components for eyes open, psilocybin: 3±1; eyes closed, psilocybin: 2.4±0.8; eyes open, placebo: 3±1; eyes closed, placebo: 2±1 [mean±STD]).

The logarithmic power spectral density (LPSD) in the delta (1–4 Hz), theta (4–8 Hz), alpha (8–12 Hz), beta (12–30 Hz) and gamma (30–40 Hz) bands was computed for each subject, condition and channel using a fast Fourier transform with a Hanning-tapered window (EEGLAB).

We estimated broadband signal complexity with the Lempel-Ziv lossless compression algorithm applied to binary time series obtained from a median split of the instantaneous signal envelope (obtained via Hilbert transform) after Z-score conversion (Schartner, et al., 2017; Timmermann et al., 2019; Pallavicini et al., 2021). By definition, the median split resulted in the same proportion of 1’s and 0’s across all channels, thus avoiding biases due to unbalanced sequences. The Lempel-Ziv algorithm divides the binary string into non-overlapping and unique binary substrings, with more substrings being required as signal diversity increases. The Lempel-Ziv complexity depends on the number of these substrings, which is maximal for a completely random sequence and minimal for a completely regular sequence.

### Local-Global auditory stimulation paradigm

The Local-Global consists of an auditory stimulation paradigm designed to evaluate responses to violations of local and global regularities, with the latter being considered a signature of conscious information processing. Following Bekinschtein et al. (2008), trials consisted of 5 brief sounds (50 ms each) separated by 100 ms. The first 4 sounds were identical, either high (1600 Hz, ‘H’) or low (800 Hz, ‘L’) pitched. The final sound of the sequence could be identical to the others or different, defining local standard (LLLLL, HHHHH) and local deviant (LLLLH, HHHHL) trials, respectively. Each block consisted of a global standard and a global deviant, importantly, the global deviant could be a local standard and vice versa. For each block, the global standard was repeated 4 times for habituation, then 4-7 global deviants were delivered interleaved with global standards, so that at least two global standards preceded each global deviant. Within each block, trials were randomly separated with silent intervals lasting 1350 to 1700 ms (in steps of 50 ms), and blocks were separated by 15 s. Finally, for each combination of global standard and deviant, blocks were repeated 5 times. This task is outlined in Figure 8A.

For the ERP analysis, EEG data was epoched and preprocessed following a similar procedure to that outlined in the “EEG” subsection. Data was bandpassed filtered in the 0.5 and 20 Hz range and baseline-corrected over a 200 ms window before the onset of each stimulus, next, ERPs (P300) corresponding to local and global deviants were constructed for the drug and the placebo conditions.

### Physical activity

The Fitbit Charge 4 wristband was used to determine two measures of physical activity during each day of the experiment: the total displacement (in kilometers) and the number of steps taken during the day. Fitbit’s tracking of step counts was shown to be accurate by direct comparison with estimates based on video recordings (Diaz et al., 2015). A systematic review showed that several measurements provided by the wristband lack sufficient accuracy; however, the step count and distance travelled were among the most accurate (Feehan et al., 2018).

### Statistical analyses

Results from both conditions (active dose and placebo) were compared using non-parametric paired Wilcoxon signed-rank tests. Unless explicitly stated, thresholds for statistical significance were Bonferroni-corrected for multiple comparisons.

This manuscript reports the analysis of all measurements obtained in our study, with the exception of EEG acquired during perception (binocular rivalry, backward masking) and cognitive tasks (attentional blink, Go / No Go, Stroop) since significant results were not found in any of these tasks at the behavioral level (see “Results” section). Also, we did not estimate statistical power when designing the experiment since we did not know *a priori* what effect size could be expected for the tasks and conditions of the experiment.

### Chemical characterization of samples

Each participant consumed 1 g of dried *Psilocybe cubensis* for the active condition, separated in two doses of 0.5 g. Samples were dried at ≈28° and ground into a fine powder, with different parts of the mushrooms homogeneously distributed (these preparations were performed by participants with the assistance of the organizers of workshops on microdosing and psilocybe mushroom home cultivation). In total, there were three independent sources for the mushrooms that were consumed in the context of this experiment. Samples (150 mg) were collected from all these sources and sent for chemical analysis to the Laboratory of Forensic Analysis of Biologically Active Substances, University of Chemistry and Technology Prague, Czech Republic. The quantification of alkaloids (psilocybin, psilocin, baeocystin and norbaeocystin) in the samples was performed by MK and KH using liquid chromatography–mass spectrometry (LC-MS). See the supplementary information for full details on the analytical methodology and chemical analysis of the samples. The concentration of the following alkaloids was determined: psilocybin (640.2 μg/g), psilocin (950.7 μg/g), baeocystin (50.4 μg/g), norbaeocystin (12.5 μg/g). At the time the chemical analysis was conducted, a single 0.5 g dose contained 0.32 mg of psilocybin and 0.48 mg of psilocin; however, at least one month passed since the end of the study and the alkaloid quantification. In spite of storing the samples under optimal conditions, the possibility that psychoactive material was lost during this period cannot be discarded (Gotvaldová et al., 2021), suggesting that the potency of the samples here reported is likely underestimated.

### Data availability

The dataset used in this manuscript can be obtained from https://zenodo.org/record/5745892

## Results

### Acute effects

We first computed the sum of all items in the VAS to obtain an index of the overall intensity of the acute effects, separately for instances when subjects correctly identified the active dose/placebo (“unblinded”) and those when this did not happen (“blinded”). Participants correctly identified dose vs. placebo in 49 of the 64 measurement weeks (75%). The VAS total scores are plotted for Wednesday (first dosing day), Thursday, Friday (second dosing day) and Saturday (last day of the experiment) in Figure 2A for the “unblinded” and “blinded” data subsets. During both dosing days the participants reported VAS total scores significantly higher for the active dose compared to the placebo, however, this difference was not present on the days when a dose was not consumed, suggesting the absence of carry-over effects. This result was also found in the “unblinded” subset of the experimental data (Figure 2A, left), but it was absent in the “blinded” one (Figure 2A, right). Figure 2B presents the outcome of this analysis conducted separately for each VAS item. While significant differences were found for items related to imagination, the dreamlike quality of the experience, spatial distortions and mind-wandering, these differences did not remain significant after applying the Bonferroni correction for multiple comparisons. Thus, the overall increase in the VAS total score depended on different, subject-specific items of the scale, instead of showing a consistent pattern across all subjects.

**Figure 2.**
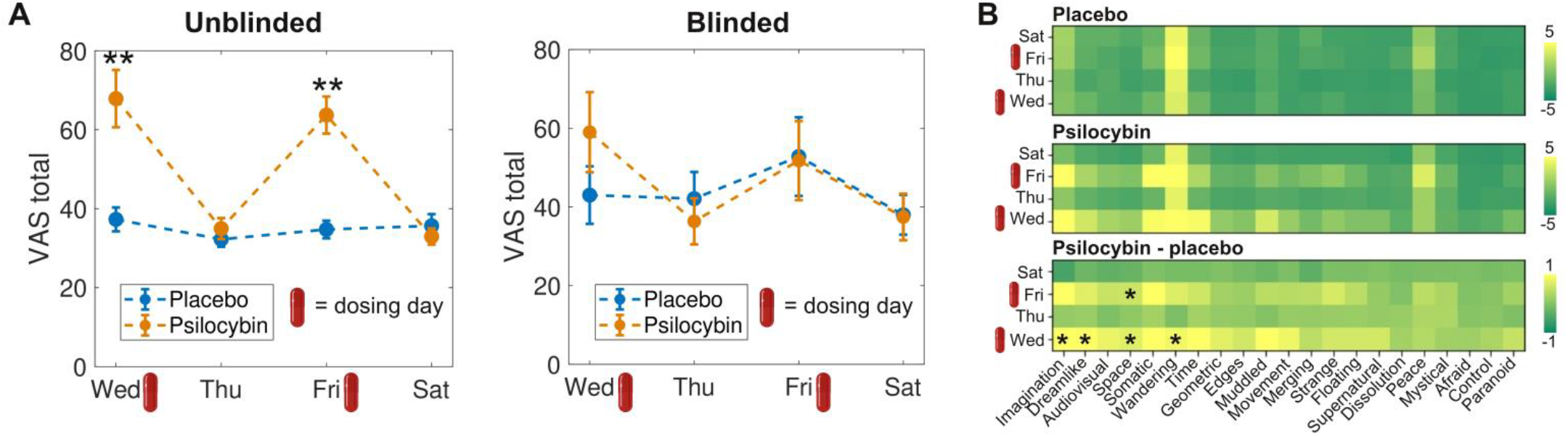
Acute effects assessed with the VAS. A) VAS total score per condition, from Wednesday (first dosing day) to Saturday (last day of the experiment), for the “unblinded” (left) and “blinded” (right) data subsets. **p<0.05, Bonferroni corrected (n=4). B) VAS scores per item, day of the experiment and experimental condition. The bottom matrix contains the difference between the active dose and the placebo. *p<0.05, uncorrected for multiple comparisons. Panel A shows mean±SEM. The small red capsules indicate the dosing days.

### Self-reported scales and questionnaires

We scored the psychometric scales and questionnaires included in Table 1, summarizing the results in Table 2, with separate columns for the “unblinded” and “blinded” data points. There were no significant differences between conditions at p<0.05, uncorrected. Figure 3 presents a boxplot of these scores for the complete dataset, with the exception of the trait variables that were only assessed at baseline (i.e. BFI, STAI-T and SSS). Note that questionnaire scores were divided by their maximum possible value and multiplied by 10, yielding normalized scores which facilitate their direct visual comparison.

**Table 2.** Results for self-reported scales and questionnaires (mean±SEM).

| Scale - Factor | Unblinded |  | Blinded |  |
| --- | --- | --- | --- | --- |
| | Mean ± SEM<br>Psilocybin | Mean ± SEM<br>Placebo | Mean ± SEM<br>Psilocybin | Mean ± SEM<br>Placebo |
| BFI - Extraversion | 29 ± 0.93 | 29.04 ± 0.99 | 30.55 ± 1.35 | 30.20 ± 1.57 |
| BFI - Agreeableness | 34.64 ± 0.81 | 35.04 ± 1.03 | 36.11 ± 1.33 | 35 ± 1.42 |
| BFI - Conscientiousness | 29.04 ± 1.01 | 28.62 ± 0.94 | 29.88 ± 1.11 | 27 ± 2.12 |
| BFI - Neuroticism | 20.37 ± 1.17 | 19.92 ± 1.25 | 18.77 ± 1.27 | 17.80 ± 1.78 |
| BFI - Openness | 42.16 ± 0.91 | 42.92 ± 1.06 | 40.11 ± 1.95 | 42.20 ± 2.20 |
| STAI-T | 3.33 ± 0.19 | 3.52 ± 0.19 | 3.40 ± 0.25 | 3.35 ± 0.42 |
| STAI-S | 2.09 ± 0.24 | 1.92 ± 0.23 | 2.22 ± 0.35 | 1.88 ± 0.32 |
| SSS | 3.34 ± 0.23 | 3.10 ± 0.17 | 3.34 ± 0.34 | 2.80 ± 0.36 |
| PANAS- | 1.36 ± 0.24 | 1.14 ± 0.22 | 1.13 ± 0.27 | 1.40 ± 0.38 |
| PANAS+ | 6.03 ± 0.37 | 5.63 ± 0.31 | 6.22 ± 0.55 | 5.55 ± 0.56 |
| PSS | 3.60 ± 0.24 | 3.34 ± 0.26 | 3.55 ± 0.44 | 3.85 ± 0.47 |
| TAS | 5.78 ± 0.40 | 5.42 ± 0.26 | 5.65 ± 0.78 | 5.52 ± 0.79 |
| BIEPS | 8.72 ± 0.13 | 8.62 ± 0.23 | 8.37 ± 0.45 | 8.30 ± 0.34 |
| MWQ | 5.36 ± 0.38 | 4.95 ± 0.32 | 5.02 ± 0.72 | 5.80 ± 0.72 |
| FSS | 5.97 ± 0.29 | 5.73 ± 0.20 | 5.81 ± 0.28 | 5.61 ± 0.37 |
| CPS | 7.16 ± 0.29 | 7.22 ± 0.24 | 7.32 ± 0.65 | 7.30 ± 0.60 |
| TECA | 5.54 ± 0.11 | 5.36 ± 0.09 | 5.53 ± 0.19 | 5.82 ± 0.21 |
| CFS | 7.96 ± 0.18 | 7.83 ± 0.21 | 7.61 ± 0.46 | 7.73 ± 0.29 |

**Figure 3.**
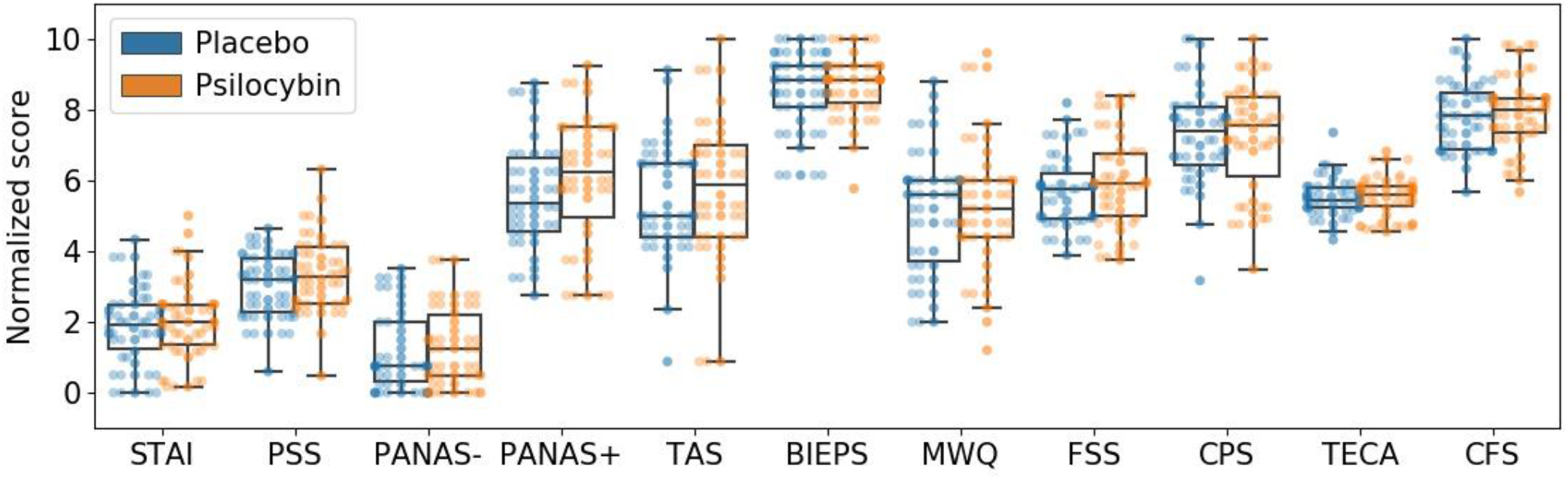
Results of the self-reported scales and questionnaires measuring state variables. Each boxplot extends from the lower to upper quartile values with a line at the median; the whiskers extend from the upper/lower quartiles up to 1.5 times the interquartile range. Subjects are represented with single points scattered on top of the boxplots. No significant differences were found at p<0.05, uncorrected.

### Creativity tests

The implemented tests were designed to measure both convergent and divergent creativity. Also, each test yielded multiple scores, each reflecting a different aspect of the performance. To measure convergent creativity, we used the RAT, scoring the total number of correct answers and the total elapsed time. To measure divergent creativity, we used the WK creativity test and the AUT. For the first, we scored the fluency (total number of words provided), the originality (responses that were given by only 5% of the group were considered unusual and scored only 1 point, responses that were given by only 1% of the group were considered unique and scored 2 points, other responses scored 0 points) and the elaboration, related to the amount of detail provided in each answer. The AUT was scored following these criteria, plus the number of repetitions (e.g. using something as a paperweight and as a way to keep papers from flying away are considered the same use). For the WK test and the AUT, higher scores are indicative of more creative performance (except for the repetition score). To avoid confounds due to different levels of fluency, the originality, repetitions and elaboration scores were normalized by the fluency score.

The results of this analysis are provided in Figure 4. There were no significant differences between conditions at p<0.05, uncorrected. Moreover, significant differences were not found when this analysis was restricted to the “unblinded” subset of data (Table 3).

**Figure 4.**
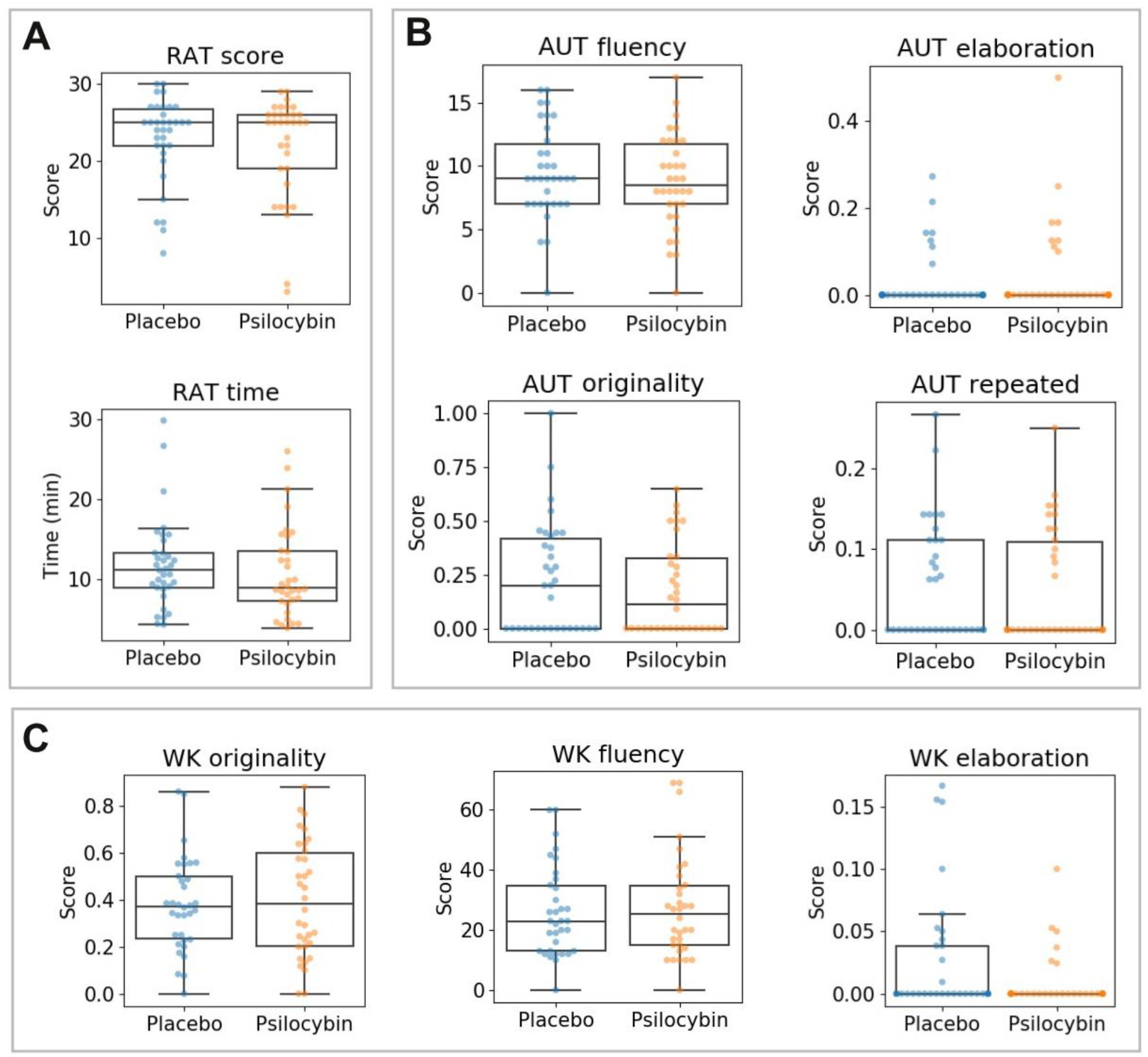
Results of creativity tests (RAT, AUY, WK). A) RAT total score and time (in minutes). B) AUT fluency, originality, elaboration and repeated answers. C) WK originality, fluency and elaboration. All boxplots extend from the lower to upper quartile values with a line at the median; the whiskers extend from the upper/lower quartiles up to 1.5 times the interquartile range. Subjects are represented with single points scattered on top of the boxplots. No significant differences were found at p<0.05, uncorrected. RAT: Remote Associations Test, AUT: Alternate Uses Test, WK: Wallach-Kogan creativity test.

**Table 3.** Results of creativity tests: RAT, AUY, and WK (mean±SEM). RAT: Remote Associations Test, AUT: Alternate Uses Test, WK: Wallace-Kogan creativity test.

| Test - Score | Unblinded |  | Blinded |  |
| --- | --- | --- | --- | --- |
| | Mean ± SEM<br>Psilocybin | Mean ± SEM<br>Placebo | Mean ± SEM<br>Psilocybin | Mean ± SEM<br>Placebo |
| RAT - Correct | 22.4±1.38 | 23.20±1.09 | 20.44±1.97 | 22.30±1.95 |
| RAT- Time (s) | 679.52±74.03 | 741.62±74.82 | 575.11±75.04 | 653.60±80.39 |
| AUT - Fluency | 8.64±0.74 | 8.79±0.70 | 9.11±1.27 | 11.30±1.17 |
| AUT - Originality | 0.16±0.03 | 0.17±0.04 | 0.23±0.08 | 0.35±0.09 |
| AUT - Repetitions | 0.04±0.01 | 0.05±0.01 | 0.06±0.02 | 0.06±0.01 |
| AUT - Elaboration | 0.05±0.02 | 0.03±0.01 | 0.01±0.01 | 0.03±0.02 |
| WK - Fluency | 26.64±3.70 | 26.79±3.32 | 29.88±4.77 | 23.90±3.59 |
| WK - Originality | 0.39±0.05 | 0.36±0.02 | 0.39±0.03 | 0.07±0.02 |
| WK - Elaboration | 0.03±0.01 | 0.03±0.01 | 0±0 | 0.02±0.01 |

### Perception tasks

Next, we investigated if microdosing with psilocybin mushrooms modulated conscious visual perception using the backward masking and the binocular rivalry tasks. The results are summarized in Figure 5, which is divided into two columns (one per task). The first panel of each column illustrates the different screens shown to the participants, together with their duration (see Materials and methods).

**Figure 5.**
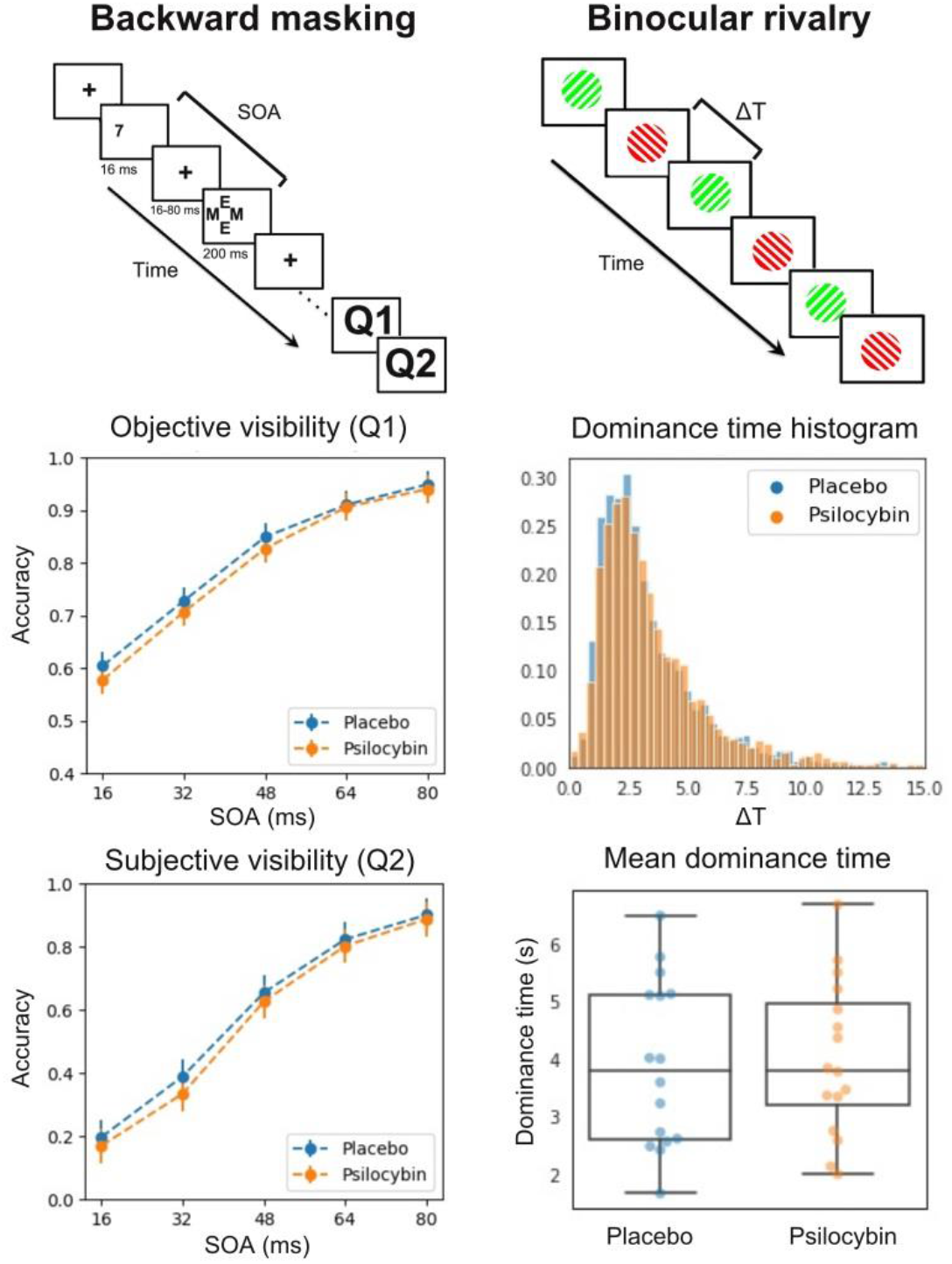
Results of the visual perception tasks. The first panel of each column contains a diagram of the corresponding task. The other two panels of the backward masking task display the accuracy vs. SOA curves for objective and subjective target visibility, respectively. The other two panels of the binocular rivalry task contain a histogram of dominance times (ΔT) pooled across subjects and separated by condition, and a boxplot of the mean dominance time per participant. The boxplot extends from the lower to upper quartile values with a line at the median; the whiskers extend from the upper/lower quartiles up to 1.5 times the interquartile range. Subjects are represented with single points scattered on top of the boxplots. SOA: stimulus onset asynchrony.

For the backward masking task, the next two panels display the accuracy vs. SOA curves for objective and subjective target visibility, respectively. As expected, the subjective visibility curve is shaped like a sigmoid, with a sharp transition in the accuracy for SOAs between 32 and 48 ms. No significant differences were found in the comparison of accuracies between conditions.

For the binocular rivalry task, we first pooled the dominance durations ΔT for both conditions across all subjects and then plotted the histograms shown in the second panel of the “Binocular rivalry” column in Figure 5. We repeated this analysis at the single subject level, fitting a gamma distribution to the histogram of individual participants (Brascamp et al., 2015). From this fit we obtained the mean dominance time per subject; these are shown in the last panel of the “Binocular rivalry” column in Figure 5. The mean dominance time is around 4 s for both conditions, without statistically significant difference between them.

### Cognitive function

Figure 6 mirrors the organization of Figure 5 but for a different set of tasks. In all cases, the first panel serves to illustrate the corresponding task (see Materials and methods). For the attentional blink, the next two panels present the visibility rate of the first (T1) and second (T2) target as a function of the lag between them (note how the accuracy drops for shorter lags, as expected). Psilocybin reduced T2 visibility for the third shortest lag compared to the placebo (results for lag 100 ms are not shown in the figure), but this difference did not hold after Bonferroni correction.

**Figure 6.**
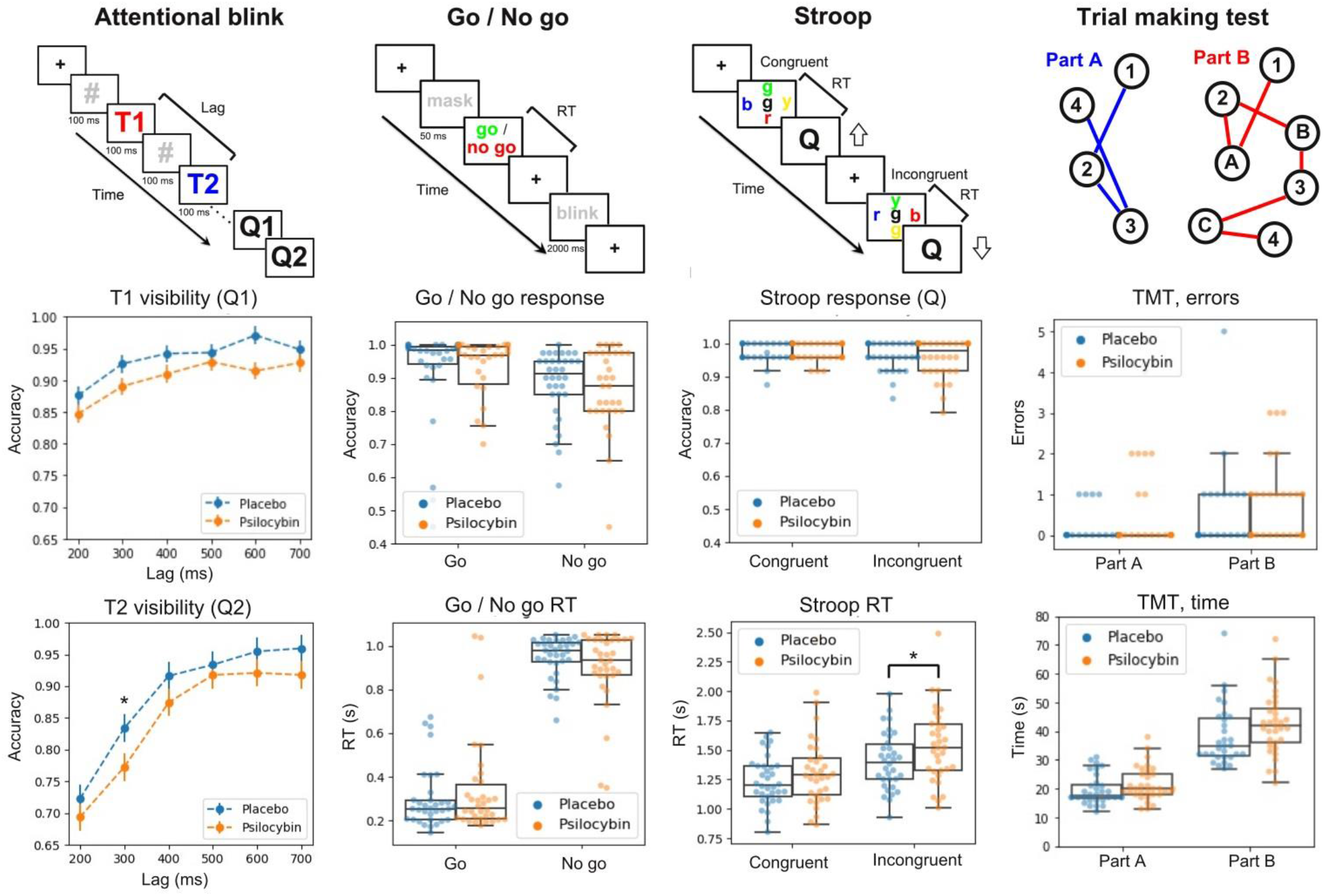
Results of tasks used to assess cognitive function. The first panel of each column contains a diagram of the task. For the attentional blink, the other two panels show the visibility of the first (T1) and second (T2) numerical target. Psilocybin reduced T2 visibility of the third shortest lag relative to placebo, but this difference did not hold after Bonferroni correction. *p<0.05 for the comparison between placebo and active dose, without correction for multiple comparisons. For the Go / No Go task, the panel shows the (normalized) number of correct responses and the reaction time. The same information is provided for the Stroop test in the third column, with a significant effect of condition on the RT of incongruent trials. *p<0.05 for the comparison between placebo and active dose, without correction for multiple comparisons. Finally, the last two panels of Figure 5 show the results of the trail making test (number of errors and time required to complete the task). All boxplots extend from the lower to upper quartile values with a line at the median; the whiskers extend from the upper/lower quartiles up to 1.5 times the interquartile range. Subjects are represented with single points scattered on top of the boxplots. TMT: trail making test, RT: reaction time.

**Figure 7.**
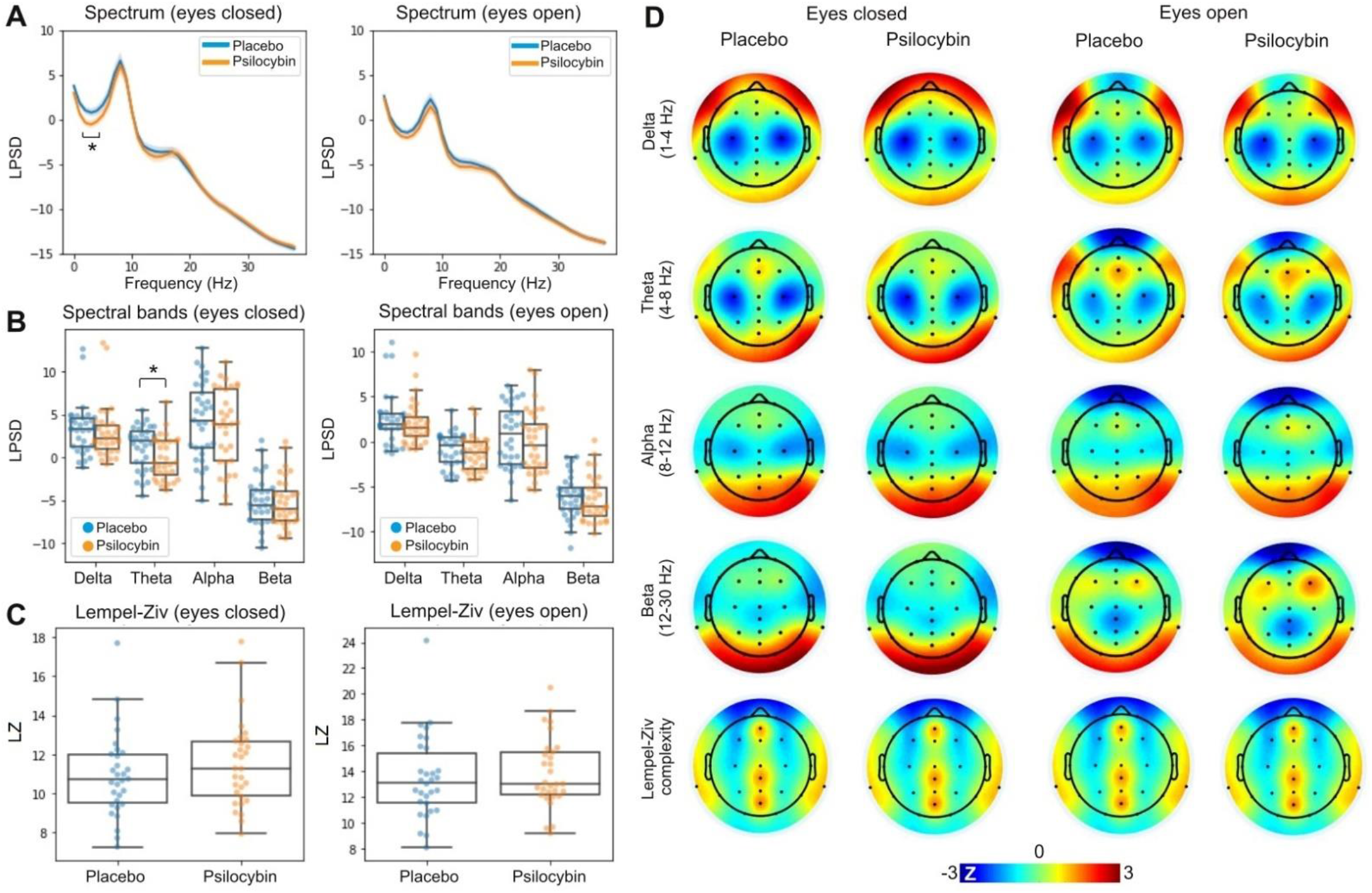
Results of the resting state EEG analysis. A) LPSD vs. frequency (averaged across all channels) for eyes closed (left) and open (right), both for the placebo and the active dose (mean±SEM). B) Same as in panel A, but binned using the following bands: delta (1-4 Hz), theta (4-8 Hz), alpha (8-12 Hz) and beta (12-20 Hz). C) Global Lempel-Ziv complexity computed using the broadband EEG signal, compared between psilocybin and placebo conditions for eyes closed and open. D) Topographic distribution of spectral power and Lempel-Ziv complexity for all combinations of active dose, placebo, eyes open and closed. All boxplots extend from the lower to upper quartile values with a line at the median; the whiskers extend from the upper/lower quartiles up to 1.5 times the interquartile range. Subjects are represented with single points scattered on top of the boxplots. *p<0.05 for the comparison between placebo and active dose. LPSD: logarithmic power spectral density.

For Go / No Go, the second and third panels in the column present boxplots of two important variables associated with this task: the response accuracy and the RT. No significant differences were found.

The last two panels on the Stroop column show the same information (response accuracy and RT). Psilocybin increased the RT of the incongruent trials relative to the placebo, but this difference did not hold after Bonferroni correction (n=4).

The last column contains the results of the trail making test; in particular, boxplots showing the total errors and the time required to complete the task.

### Resting state EEG

Figure 6 presents the results of the resting state EEG analysis. Panel A displays the LPSD vs. frequency (averaged across all channels) for eyes closed (left) and open (right), both for the placebo and the active dose. As expected, the power spectra for eyes closed show a peak close to 10 Hz, which is attenuated for the eyes open condition. The psilocybin mushroom microdose resulted in decreased theta power in the theta range (4 to 8 Hz). Figure 6B presents the same results binned into four major frequency bands: delta (1-4 Hz), theta (4-8 Hz), alpha (8-12 Hz) and beta (12-20 Hz). Consistent with the spectra shown in Figure 6A, only the eyes closed theta band power decreased under the active dose compared to the placebo. Figure 6C compares the global Lempel-Ziv complexity computed using the broadband EEG signal between psilocybin and placebo conditions for eyes closed (left) and open (right). Finally, Figure 6D presents the topographic distribution of spectral power and Lempel-Ziv complexity for all combinations of active dose, placebo, eyes open and closed.

### Local-global evoked response potentials

We computed the ERPs associated with local and global deviants from the Local-Global auditory stimulation. Figure 8A outlines the structure of trials and blocks in the task (see Materials and methods). For all channels, we investigated whether the global deviant resulted in larger late amplitude deflections compared to the local deviant, as expected from the previous literature (Bekinschtein et al., 2008). Also consistent with previous work, the central-frontal channel AFz presented the largest effects. Figure 8B shows the global deviant minus the local deviant ERPs located at AFz for the placebo condition; clearly, after 300 ms the global deviant presents a more sustained amplitude, which has been linked to conscious information processing. Panels C and D show the ERPs at AFz for local and global deviants, respectively, and for the active dose and placebo conditions. No significant differences between conditions were found.

**Figure 8.**
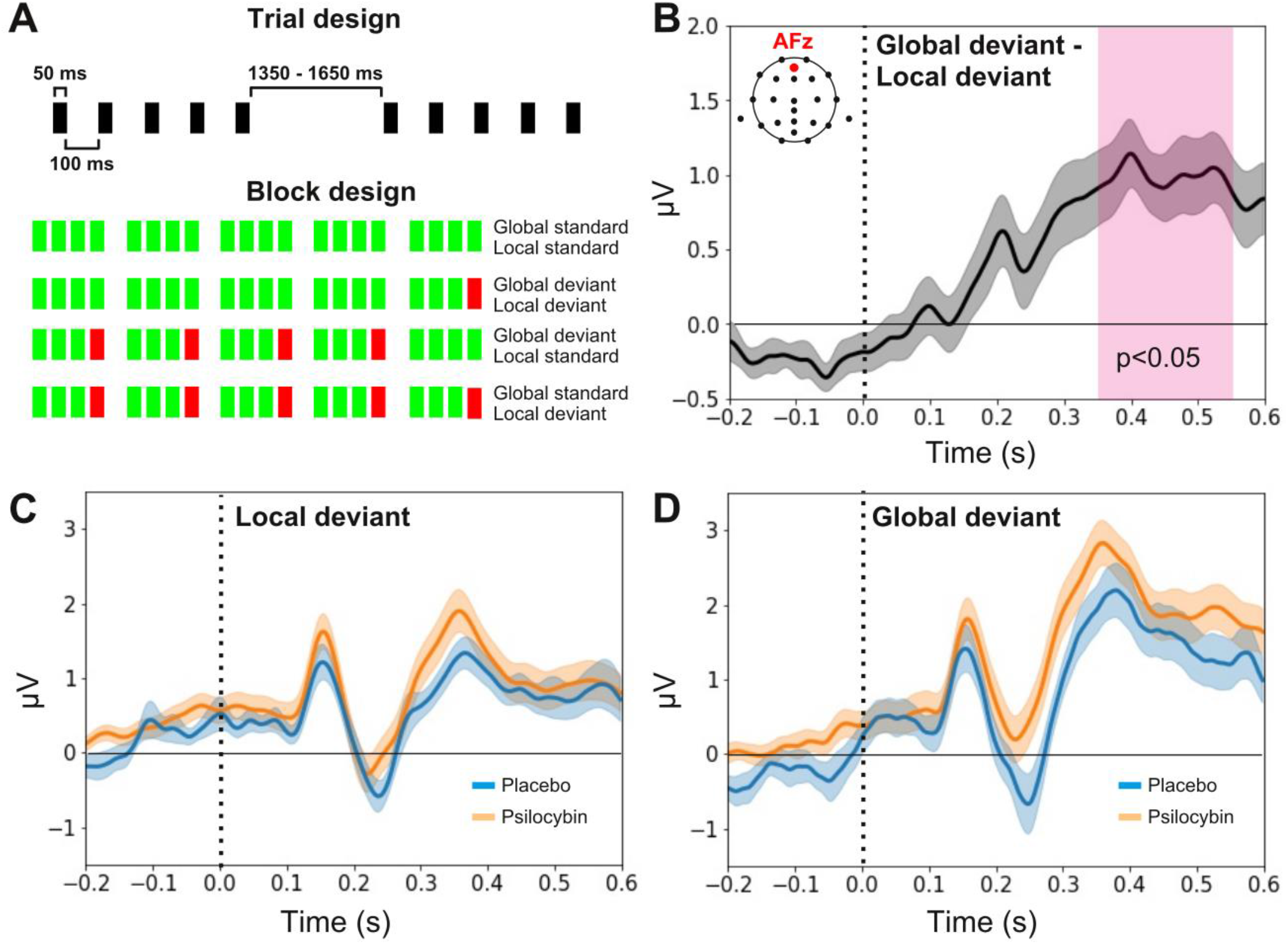
Results of the Local-Global ERP analysis. A) Structure of trials and blocks in the task (see Materials and methods). B) Global deviant minus the local deviant ERPs located at AFz for the placebo condition. C) Local deviant at AFz for placebo vs. psilocybin. D) Same as in panel C but for the global deviant. All ERP plots show mean±SEM. The vertical dashed lines coincide with the timing of the last sound in the trial.

### Physical activity

Figure 9 summarizes the results of the physical activity analysis based on data provided by the Fitbit Charger 4 wristband. Figures 9A to 9D present the step count, distance traveled, resting and activity time, respectively, per each day of the week during the experiment. We did not find differences between the active dose and placebo conditions. The decrease in activity on Wednesday that is seen for step count, distance travelled and activity time may occur because several measurements that require sitting to complete tasks were conducted on that day.

**Figure 9.**
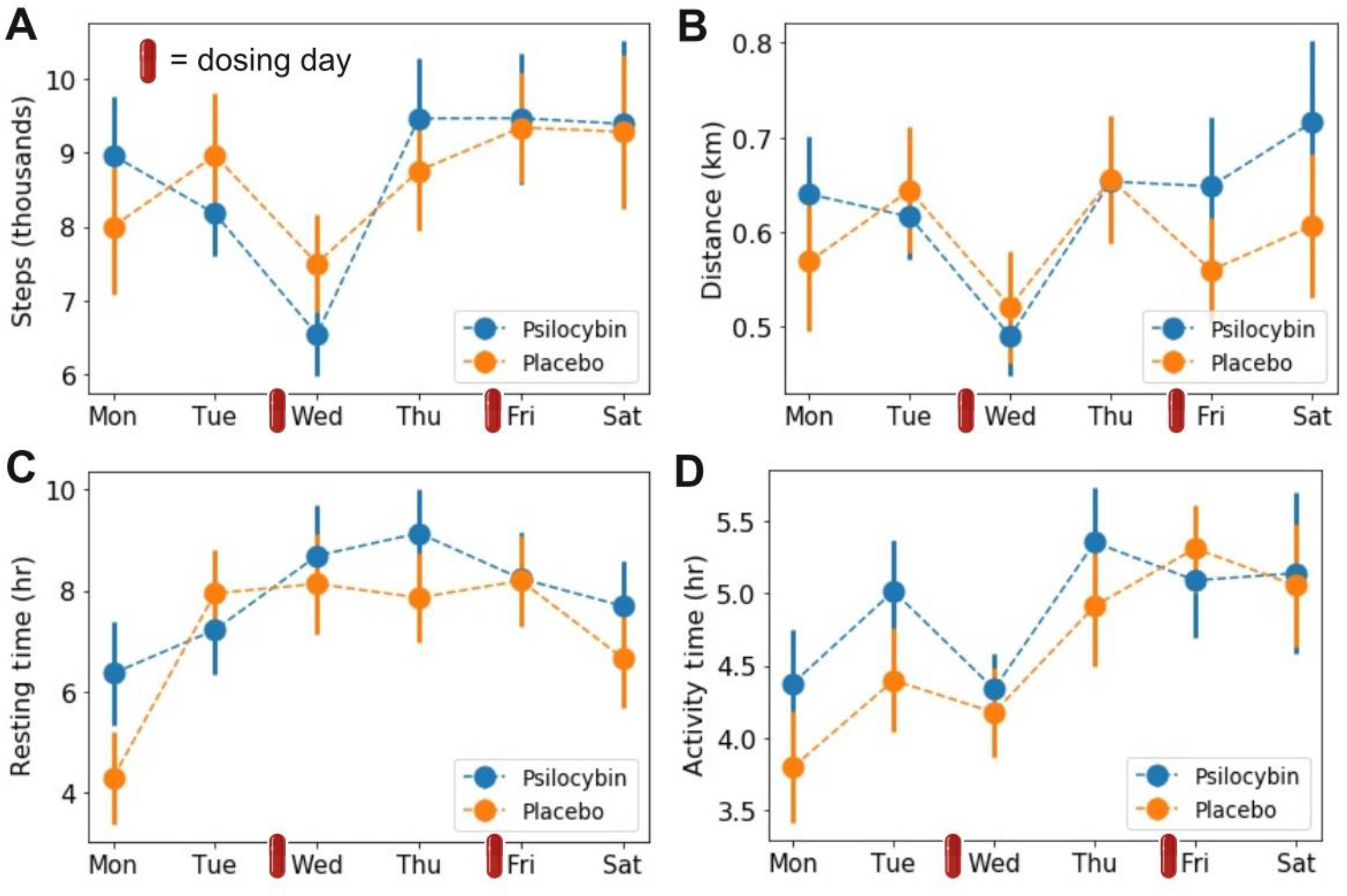
Results of the physical activity analysis. A) Step count per week day. B) Distance travelled per week day. C) Resting time per week day. D) Total activity time per week day. All plots show mean±SEM. The small red capsules indicate the dosing days.

### Results for the unblinded group

We repeated the statistical analyses for the results shown in Figures 5 to 9, but dividing them into “unblinded” and “blinded” subsets. We expected that unblinding would result in larger differences between conditions due to confirmation bias, as seen in previous microdosing studies, where many of the reported significant effects were attributed to breaking of the blinding. However, this analysis did not yield statistically significant results.

## Discussion

We investigated whether microdosing with *Psilocybe cubensis* mushrooms affected different aspects of human behavior, cognition, perception, as well as divergent and convergent thinking. This was complemented with an assessment of several self-reported scales and questionnaires used to measure personality, anxiety, mood, well-being, absorption, positive and negative thinking, mind-wandering, flow, and cognitive flexibility, among other relevant constructs. Our protocol also included one of the first measurements of electroencephalography and physical activity within the context of microdosing with psychedelic drugs. According to our results, 0.5 g of dried mushroom material did not significantly impact in any of these domains, although we observed a trend towards impaired performance in some cognitive tasks (attentional blink and Stroop). In contrast, the overall acute effects induced by the microdose (VAS total score) were significant, although they lacked consistency across participants. We also found decreased EEG power in the theta band under psilocybin, which is consistent with the broadband spectral power reductions reported for higher doses.

Ample anecdotal evidence suggests that microdosing can improve mood, well-being, creativity, and cognition (Fadiman, 2011; Fadiman and Krob, 2017), and recent uncontrolled, open-label observational studies have provided some empirical support for these claims (Johnstad, 2018; Prochazkova et al., 2018; Hutten et al., 2019; Polito and Stevenson, 2019; Anderson et al., 2019a; Anderson et al., 2019b; Fadiman and Korb, 2019; Webb et al., 2019; Cameron et al., 2020; Lea et al., 2020). While encouraging, these studies are vulnerable to experimental biases, including confirmation bias and placebo effects (Muthukumaraswamy et al., 2021). This is especially problematic in the case of microdosing, since users make up a self-selected sample with optimistic expectations about the outcome of the practice (Polito and Stevenson, 2019; Ona and Bousa, 2020). This positivity bias, combined with the low doses and self-assessment of the drug effects via scales and questionnaires, paves the way for a strong placebo response (Muthukumaraswamy et al., 2021).

To date, we could identify relatively few human studies of microdosing with psychedelics following a rigorous experimental design. The first was conducted by Yanakieva and colleagues, who investigated three comparatively low doses of LSD (5, 10, and 20 μg) (Yanakieva et al., 2019), concluding that LSD affected the estimation of time intervals, without other significant changes in perception, mental processes and concentration. However, the researchers did not assess the preexisting motivations and expectations of the participants, and the laboratory setting of the experiment might have contributed to their suboptimal performance. Bershad and colleagues investigated an inactive placebo and three different doses of LSD (6.3, 13, and 26 μg) separated by one-week intervals (Bershad et al., 2019). At the highest dose, the drug increased ratings of vigor and slightly decreased positivity ratings of images with positive emotional content. Measurements of mood, cognition, and physiological responses did not show differences between conditions. Another study by the same group (Bershad et al., 2020) showed that a low dose of LSD (13 μg) increased amygdala seed-based connectivity with the right angular gyrus, right middle frontal gyrus, and the cerebellum, and decreased amygdala connectivity with the left and right postcentral gyrus and the superior temporal gyrus. Although this dose of LSD had weak effects on mood, they were positively correlated with the increase in amygdala–middle frontal gyrus connectivity strength. Family et al., (2020) established the safety of LSD microdosing in older volunteers, but did not report substantial positive effects. Hutten and colleagues reported dose-dependent positive effects on mood, but also anxiety and cognitive impairment (Hutten et al., 2020); also, the same group showed that low doses of LSD can increase brain-derived neurotrophic factor blood plasma levels in healthy volunteers (Hutten et al., 2021). Finally, both Szigeti et al. (2021) and Van Elk et al. (2021) combined double-blind placebo-controlled design with field measurements under natural conditions. Both found a positive effect of microdosing on the primary outcome of their respective studies; however, these results could be explained by breaking of the placebo condition. In particular, Van Elk et al. and Bershad et al. found that more than 60% and 80% of the participants were breaking blind to their experimental condition, respectively (Bershad et al., 2020; Van Elk et al., 2021), consistently with the unblinding rate found in our study (75%).

Our results add to this series of double-blind placebo-controlled studies questioning the validity of anecdotal evidence for microdosing (Ona and Bouso, 2020). In comparison to previous studies (Bershad et al., 2020; Van Elk et al., 2021), most results remained negative even when the statistical analyses were restricted to measurements obtained from unblinded subjects (with the exception of the VAS total scores of acute effects, see Figure 2A). Overall, few uncorrected differences were found, in all cases indicative of impaired performance, which is consistent with previous experiments (Hutten et al., 2020) and with the observation that higher doses of serotonergic psychedelics negatively affect cognitive functions such as attention and decision making (Bayne and Carter, 2018). It has also been suggested that psychedelics might facilitate visual perception by increasing the broadband of consciously perceived information (Bayne and Carter, 2018). This was supported by studies of binocular rivalry, showing that two doses of psilocybin (115 μg/kg and 250 μg/kg) slowed down the rate of binocular rivalry switching and increased the proportion of reports of mixed percepts. Our study failed to replicate these findings, possibly due to the lower effective dose of psilocybin contained in the mushroom preparations. Also, we directly investigated the potential influence of microdosing on conscious perception using a backward masking paradigm (for visual perception, Del Cul et al., 2007) and the Global-Local paradigm combined with EEG for ERP analysis of global and local deviants (for auditory perception; Bekinschtein et al., 2008). Neither of these tasks revealed a significant effect of *Psilocybe cubensis* microdosing on conscious information processing.

One of the most robust neurophysiological markers of psychedelic action is given by scalp EEG, a technique that consistently demonstrated reduced oscillatory power as a consequence of 5-HT_2A_ receptor activation (Riba et al., 2002; Muthukumaraswamy et al., 2013; Kometer et al., 2013; Schenberg et al., 2015; Valle et al., 2016; Carhart-Harris et al., 2016; Tagliazucchi et al., 2021). More recently, spectral power analyses began to be complemented with the investigation of signal diversity, for instance, by means of the Lempel-Ziv complexity (Schartner et al., 2017; Timmermann et al., 2019; Pallavicini et al., 2021). In line with the entropic brain hypothesis (Carhart-Harris et al., 2014), EEG signal complexity has been shown to increase during the acute effects of several serotonergic psychedelics (LSD, psilocybin, DMT) and ketamine, a glutamatergic dissociative (Schartner et al., 2017). We found reduced power of theta oscillations during the effects of the psilocybin microdose, heralding the larger broadband reductions observed for higher doses (Murray et al., 2021). However, the analysis of Lempel-Ziv complexity failed to reveal differences between conditions, suggesting that increased signal entropy might constitute a specific signature of the altered consciousness elicited by psychedelics or other non-pharmacological mechanisms (Vivot et al., 2020; Farnes et al., 2020; Aamodt et al., 2021).

We investigated daily levels of physical activity as a proxy of the potential effects of microdosing on mood and well-being. The relationship between physical activity and mental health is well-established (Paluska and Schwenk, 2000), and has been adopted as a marker of treatment efficacy for depression (Roshanaei-Moghaddam et al., 2009). Currently, the potential association between changes in physical activity levels and psychedelic use remains unexplored. While our results did not reveal an effect of microdosing on this domain, future studies could further this investigation using higher doses of serotonergic psychedelics, both in healthy and clinical populations (Kuypers et al., 2019), and conducting measurements over longer time periods.

While the study of microdosing with *Psilocybe cubensis* mushrooms presents advantages in terms of ecological validity, it also raises problems associated with unknown or inconsistent chemical composition. We analyzed the contents of three samples pooled together, estimating an effective dose of ≈0.9 mg of psilocybin; however, this dose could have been higher or lower depending on the source of the mushrooms consumed by each participant. Also, we did not correct the effective psilocybin dose using the weight of the participants. While this adjustment might not be necessary for larger doses, its importance for microdosing remains unexplored (Garcia-Romeu et al., 2021). The amount of psilocybin/psilocin found in our samples is well within the expected values for the mushrooms or truffles that are consumed in the context of microdosing (Polito and Stevenson, 2019; Van Elk et al., 2021; Szigeti et al., 2021); in particular, it is almost identical to the values reported in Prochazkova et al. (2018). Nevertheless, other recent studies used truffles with higher concentrations of psilocin and psilocybin; for instance, Van Elk and colleagues investigated the effects of 0.7 g of psilocybin-containing truffles, with an estimated amount of 1.5 mg of psilocybin per dose. As acknowledged by the authors of this particular study, 0.7 g exceeds what is frequently considered the upper limit when microdosing with psilocybin mushrooms (note, however, that what constitutes “microdosing” is not precisely defined). It is also important to consider the possibility that our samples lost potency between the experiment and their chemical analysis. As shown by Gotvaldová and colleagues, the concentration of psilocybin can drop up to 50% during the first months of storage (Gotvaldová et al., 2021), which in our case would imply original concentrations similar to those reported by Van Elk and colleagues. Finally, our samples contained small amounts baeocystin and norbaeocystin; whether these compounds are psychoactive in humans is still under discussion (Gartz, 1992; Sherwood et al., 2020).

Microdosing is generally conducted over extended periods of time according to different dosing schedules (Fadiman, 2011). By design, our study could not assess the cumulative effects of microdoses consumed over periods of several days. Due to the known build-up of tolerance after repeated administration of serotonergic psychedelics (Nichols, 2016), we speculated that the intensity of these effects could only decrease in time; because of this, we decided to investigate the acute effects of microdosing instead of its potential cumulative effects. Future research should explore whether the positive effects of microdosing can be selectively enabled or facilitated by certain long-term dosing schedules.

In conclusion, we conducted a controlled study of microdosing in individuals who were already planning to start their own microdosing protocol. While small amounts of dried *Psilocybe cubensis* mushrooms reliably induced significant subjective effects, their impact in other domains was negligible or even indicative of impaired performance. Clearly, more research is needed to decide whether microdosing with psychedelics can deliver at least some of its promised positive effects. This future research should also explore the potential impact of microdosing on aspects of human physiology that could compromise its long-term safety; for instance, by addressing the potential consequences of chronic 5-HT_2A_ receptor stimulation on the health of the circulatory system, among other important points (Kuypers et al., 2019; Ona and Bouso, 2020). Until this research is conducted, it remains impossible to ascertain that microdosing is a safe practice leading to desirable effects, and to rule out that these effects arise as a consequence of expectation or confirmation biases.

## Acknowledgments

We thank Sol Pérez Vázquez for her assistance with the logistics of this study. This study was funded by grant PICT-2019-02294 awarded by Agencia Nacional de Promoción Científica y Tecnológica (Argentina). Partial funding was received from the Ministry of Health of the Czech Republic (grant no. NU21-04-00307) and the Grant Agency of the Czech Republic (grant no. 20-25349S).

## References

Aamodt, A., Nilsen, A. S., Thürer, B., Moghadam, F. H., Kauppi, N., Juel, B. E., & Storm, J. F. (2021). EEG signal diversity varies with sleep stage and aspects of dream experience. Frontiers in psychology, 12, 1204.

Amabile, T. M. (1985). Motivation and creativity: Effects of motivational orientation on creative writers. Journal of personality and social psychology, 48(2), 393.

Andersson, M., Persson, M., & Kjellgren, A. (2017). Psychoactive substances as a last resort—a qualitative study of self-treatment of migraine and cluster headaches. Harm reduction journal, 14(1), 1–10.

Anderson, T., Petranker, R., Christopher, A., Rosenbaum, D., Weissman, C., Dinh-Williams, L. A.,… & Hapke, E. (2019a). Psychedelic microdosing benefits and challenges: an empirical codebook. Harm reduction journal, 16(1), 1–10.

Anderson, T., Petranker, R., Rosenbaum, D., Weissman, C. R., Dinh-Williams, L. A., Hui, K.,… & Farb, N. A. (2019b). Microdosing psychedelics: personality, mental health, and creativity differences in microdosers. Psychopharmacology, 236(2), 731–740.

Bayne, T., & Carter, O. (2018). Dimensions of consciousness and the psychedelic state. Neuroscience of consciousness, 2018(1), niy008.

Bekinschtein, T. A., Dehaene, S., Rohaut, B., Tadel, F., Cohen, L., & Naccache, L. (2009). Neural signature of the conscious processing of auditory regularities. Proceedings of the National Academy of Sciences, 106(5), 1672–1677.

Benet-Martínez, V., & John, O. P. (1998). Los Cinco Grandes across cultures and ethnic groups: Multitrait-multimethod analyses of the Big Five in Spanish and English. Journal of personality and social psychology, 75(3), 729.

Bershad, A. K., Schepers, S. T., Bremmer, M. P., Lee, R., & de Wit, H. (2019). Acute subjective and behavioral effects of microdoses of lysergic acid diethylamide in healthy human volunteers. Biological psychiatry, 86(10), 792–800.

Bershad, A. K., Preller, K. H., Lee, R., Keedy, S., Wren-Jarvis, J., Bremmer, M. P., & de Wit, H. (2020). Preliminary report on the effects of a low dose of LSD on resting-state amygdala functional connectivity. Biological Psychiatry: Cognitive Neuroscience and Neuroimaging, 5(4), 461–467.

Brascamp, J. W., Klink, P. C., & Levelt, W. J. (2015). The ‘laws’ of binocular rivalry: 50 years of Levelt’s propositions. Vision research, 109, 20–37.

Brunner, C., Delorme, A., & Makeig, S. (2013). Eeglab–an open source matlab toolbox for electrophysiological research. Biomedical Engineering/Biomedizinische Technik, 58(SI-1-Track-G), 000010151520134182.

Cameron, L. P., Nazarian, A., & Olson, D. E. (2020). Psychedelic microdosing: prevalence and subjective effects. Journal of psychoactive drugs, 52(2), 113–122.

Castro, A., Brenlla, M., & Casullo, M. (2002). Evaluación del bienestar psicológico en adultos argentinos. Evaluación del bienestar psicológico en Iberoamérica. Buenos Aires: Paidós.

Carhart-Harris, R. L., Muthukumaraswamy, S., Roseman, L., Kaelen, M., Droog, W., Murphy, K.,… & Nutt, D. J. (2016). Neural correlates of the LSD experience revealed by multimodal neuroimaging. Proceedings of the National Academy of Sciences, 113(17), 4853–4858.

Carhart-Harris, R. L., Leech, R., Hellyer, P. J., Shanahan, M., Feilding, A., Tagliazucchi, E.,… & Nutt, D. (2014). The entropic brain: a theory of conscious states informed by neuroimaging research with psychedelic drugs. Frontiers in human neuroscience, 8, 20.

Carhart-Harris, R. L., Roseman, L., Haijen, E., Erritzoe, D., Watts, R., Branchi, I., & Kaelen, M. (2018). Psychedelics and the essential importance of context. Journal of Psychopharmacology, 32(7), 725–731.

Carter, O. L., Hasler, F., Pettigrew, J. D., Wallis, G. M., Liu, G. B., & Vollenweider, F. X. (2007). Psilocybin links binocular rivalry switch rate to attention and subjective arousal levels in humans. Psychopharmacology, 195(3), 415–424.

Carter, O., Pettigrew, J., Hasler, F., Wallis, G., & Vollenweider, F. (2010). Psilocybin slows binocular rivalry switching through serotonin modulation. Journal of Vision, 6(6), 43.

Cohen, S., Kamarck, T., & Mermelstein, R. (1994). Perceived stress scale. Measuring stress: A guide for health and social scientists, 10(2), 1–2.

Del Cul, A., Baillet, S., & Dehaene, S. (2007). Brain dynamics underlying the nonlinear threshold for access to consciousness. PLoS biology, 5(10), e260.

Delorme, A., & Makeig, S. (2004). EEGLAB: an open source toolbox for analysis of single-trial EEG dynamics including independent component analysis. Journal of neuroscience methods, 134(1), 9–21.

Diaz, K. M., Krupka, D. J., Chang, M. J., Peacock, J., Ma, Y., Goldsmith, J.,… & Davidson, K. W. (2015). Fitbit®: An accurate and reliable device for wireless physical activity tracking. International journal of cardiology, 185, 138.

Fadiman, J. (2011). The psychedelic explorer’s guide: Safe, therapeutic, and sacred journeys. Simon and Schuster.

Fadiman, J., & Krob, S. (2017). Microdosing: the phenomenon, research results, and startling surprises. In Lecture presented at the Psychedelic Science 2017 conference, Oakland, CA.

Fadiman, J., & Korb, S. (2019). Might microdosing psychedelics be safe and beneficial? An initial exploration. Journal of psychoactive drugs, 51(2), 118–122.

Family, N., Maillet, E. L., Williams, L. T., Krediet, E., Carhart-Harris, R. L., Williams, T. M.,… & Raz, S. (2020). Safety, tolerability, pharmacokinetics, and pharmacodynamics of low dose lysergic acid diethylamide (LSD) in healthy older volunteers. Psychopharmacology, 237(3), 841–853.

Farnes, N., Juel, B. E., Nilsen, A. S., Romundstad, L. G., & Storm, J. F. (2020). Increased signal diversity/complexity of spontaneous EEG, but not evoked EEG responses, in ketamine-induced psychedelic state in humans. Plos one, 15(11), e0242056.

Feehan, L. M., Geldman, J., Sayre, E. C., Park, C., Ezzat, A. M., Yoo, J. Y.,… & Li, L. C. (2018). Accuracy of Fitbit devices: systematic review and narrative syntheses of quantitative data. JMIR mHealth and uHealth, 6(8), e10527.

Garcia-Romeu, A., Barrett, F. S., Carbonaro, T. M., Johnson, M. W., & Griffiths, R. R. (2021). Optimal dosing for psilocybin pharmacotherapy: Considering weight-adjusted and fixed dosing approaches. Journal of Psychopharmacology, 35(4), 353–361.

Gartz, J. (1992). Further investigations on psychoactive mushrooms of the genera Psilocybe, Gymnopilus and Conocybe. editore non identificato.

Glatter, R. (2015). LSD microdosing: The new job enhancer in Silicon Valley and beyond. Forbes.

Gotvaldová, K., Hájková, K., Borovička, J., Jurok, R., Cihlářová, P., & Kuchař, M. (2021). Stability of psilocybin and its four analogs in the biomass of the psychotropic mushroom Psilocybe cubensis. Drug testing and analysis, 13(2), 439–446.

Gough, H. G. (1979). A creative personality scale for the adjective check list. Journal of personality and social psychology, 37(8), 1398.

Guilford, J. P. (1967). The nature of human intelligence.

Hutten, N. R., Mason, N. L., Dolder, P. C., & Kuypers, K. P. (2019a). Motives and side-effects of microdosing with psychedelics among users. International Journal of Neuropsychopharmacology, 22(7), 426–434.

Hutten, N. R., Mason, N. L., Dolder, P. C., & Kuypers, K. P. (2019b). Self-rated effectiveness of microdosing with psychedelics for mental and physical health problems among microdosers. Frontiers in psychiatry, 10, 672.

Hutten, N. R., Mason, N. L., Dolder, P. C., Theunissen, E. L., Holze, F., Liechti, M. E.,… & Kuypers, K. P. (2020). Mood and cognition after administration of low LSD doses in healthy volunteers: a placebo controlled dose-effect finding study. European Neuropsychopharmacology, 41, 81–91.

Hutten, N. R., Mason, N. L., Dolder, P. C., Theunissen, E. L., Holze, F., Liechti, M. E.,… & Kuypers, K. P. (2021). Low doses of LSD acutely increase BDNF blood plasma levels in healthy volunteers. ACS Pharmacology & Translational Science, 4(2), 461–466.

Jackson, S. A., & Marsh, H. W. (1996). Development and validation of a scale to measure optimal experience: The Flow State Scale. Journal of sport and exercise psychology, 18(1), 17–35.

Johnson, M. W., Richards, W. A., & Griffiths, R. R. (2008). Human hallucinogen research: guidelines for safety. Journal of psychopharmacology, 22(6), 603–620.

Johnstad, P. G. (2018). Powerful substances in tiny amounts: an interview study of psychedelic microdosing. Nordic Studies on Alcohol and Drugs, 35(1), 39–51.

Kaertner, L. S., Steinborn, M. B., Kettner, H., Spriggs, M. J., Roseman, L., Buchborn, T.,… & Carhart-Harris, R. L. (2021). Positive expectations predict improved mental-health outcomes linked to psychedelic microdosing. Scientific reports, 11(1), 1–11.

Kometer, M., Schmidt, A., Jäncke, L., & Vollenweider, F. X. (2013). Activation of serotonin 2A receptors underlies the psilocybin-induced effects on α oscillations, N170 visual-evoked potentials, and visual hallucinations. Journal of Neuroscience, 33(25), 10544–10551.

Kotov, R. I., Bellman, S. B., & Watson, D. B. (2004). Short Suggestibility Scale.

Kuypers, K. P., Ng, L., Erritzoe, D., Knudsen, G. M., Nichols, C. D., Nichols, D. E.,… & Nutt, D. (2019). Microdosing psychedelics: More questions than answers? An overview and suggestions for future research. Journal of Psychopharmacology, 33(9), 1039–1057.

Kuypers, K. P. (2020). The therapeutic potential of microdosing psychedelics in depression. Therapeutic advances in psychopharmacology, 10, 2045125320950567.

Lea, T., Amada, N., Jungaberle, H., Schecke, H., Scherbaum, N., & Klein, M. (2020). Perceived outcomes of psychedelic microdosing as self-managed therapies for mental and substance use disorders. Psychopharmacology, 1–12.

Lea, T., Amada, N., Jungaberle, H., Schecke, H., & Klein, M. (2020). Microdosing psychedelics: motivations, subjective effects and harm reduction. International Journal of Drug Policy, 75, 102600.

López-Pérez, B., Fernández-Pinto, I., & García, F. J. A. (2008). TECA: Test de empatía cognitiva y afectiva. Tea.

Mrazek, M. D., Phillips, D. T., Franklin, M. S., Broadway, J. M., & Schooler, J. W. (2013). Young and restless: validation of the Mind-Wandering Questionnaire (MWQ) reveals disruptive impact of mind-wandering for youth. Frontiers in psychology, 4, 560.

Martin, M. M., & Rubin, R. B. (1995). A new measure of cognitive flexibility. Psychological reports, 76(2), 623–626.

Mednick, S. A., & Mednick, M.T. (1959). Remote Associates Test, college and adult form.

Muthukumaraswamy, S. D., Carhart-Harris, R. L., Moran, R. J., Brookes, M. J., Williams, T. M., Errtizoe, D.,… & Nutt, D. J. (2013). Broadband cortical desynchronization underlies the human psychedelic state. Journal of Neuroscience, 33(38), 15171–15183.

Muthukumaraswamy, S., Forsyth, A., & Lumley, T. (2021). Blinding and expectancy confounds in psychedelic randomised controlled trials. Expert Review of Clinical Pharmacology, (in press).

Murray, C. H., Tare, I., Perry, C. M., Malina, M., Lee, R., & de Wit, H. (2021). Low doses of LSD reduce broadband oscillatory power and modulate event-related potentials in healthy adults. Psychopharmacology, 1–13.

Nichols, D. E. (2016). Psychedelics. Pharmacological reviews, 68(2), 264–355.

Nye, B. C. Microdosing: The people taking LSD with their breakfast. BBC News. BBC News (2017) https://www.bbc.com/news/health-39516345.

Olson, J. A., Suissa-Rocheleau, L., Lifshitz, M., Raz, A., & Veissiere, S. P. (2020). Tripping on nothing: placebo psychedelics and contextual factors. Psychopharmacology, 237(5), 1371–1382.

Olson, D. E. (2020). The subjective effects of psychedelics may not be necessary for their enduring therapeutic effects. ACS Pharmacology & Translational Science, 4(2), 563–567.

Ona, G., & Bouso, J. C. (2020). Potential safety, benefits, and influence of the placebo effect in microdosing psychedelic drugs: A systematic review. Neuroscience & Biobehavioral Reviews.

Pallavicini, C., Cavanna, F., Zamberlan, F., de la Fuente, L. A., Ilksoy, Y., Perl, Y. S.,… & Tagliazucchi, E. (2021). Neural and subjective effects of inhaled N, N-dimethyltryptamine in natural settings. Journal of Psychopharmacology, 35(4), 406–420.

Paluska, S. A., & Schwenk, T. L. (2000). Physical activity and mental health. Sports medicine, 29(3), 167–180.

Passie, T., Seifert, J., Schneider, U., & Emrich, H. M. (2002). The pharmacology of psilocybin. Addiction biology, 7(4), 357–364.

Polito, V., & Stevenson, R. J. (2019). A systematic study of microdosing psychedelics. PloS one, 14(2), e0211023.

Prabhu, V., Sutton, C., & Sauser, W. (2008). Creativity and certain personality traits: Understanding the mediating effect of intrinsic motivation. Creativity Research Journal, 20(1), 53–66.

Prochazkova, L., Lippelt, D. P., Colzato, L. S., Kuchar, M., Sjoerds, Z., & Hommel, B. (2018). Exploring the effect of microdosing psychedelics on creativity in an open-label natural setting. Psychopharmacology, 235(12), 3401–3413.

Reitan, R. M., & Wolfson, D. (1995). Category Test and Trail Making Test as measures of frontal lobe functions. The Clinical Neuropsychologist, 9(1), 50–56.

Riba, J., Anderer, P., Morte, A., Urbano, G., Jané, F., Saletu, B., & Barbanoj, M. J. (2002). Topographic pharmaco-EEG mapping of the effects of the South American psychoactive beverage ayahuasca in healthy volunteers. British journal of clinical pharmacology, 53(6), 613–628.

Roshanaei-Moghaddam, B., Katon, W. J., & Russo, J. (2009). The longitudinal effects of depression on physical activity. General hospital psychiatry, 31(4), 306–315.

Rootman, J., Kryskow, P., Harvey, K., Stamets, P., Santos-Brault, E., Kuypers, K., Polito, V., Bourzat, F., Walsh, Z. (2021). Adults who microdose psychedelics report health related motivations and lower levels of anxiety and depression compared to non-microdosers. Scientific reports (in press).

Sahakian, B., d’Angelo, C., & Savulich, G. (2018). Microdosing’LSD Is Not Just a Silicon Valley Trend–It Is Spreading to Other Workplaces. The Independent.

Schartner, M. M., Carhart-Harris, R. L., Barrett, A. B., Seth, A. K., & Muthukumaraswamy, S. D. (2017). Increased spontaneous MEG signal diversity for psychoactive doses of ketamine, LSD and psilocybin. Scientific reports, 7(1), 1–12.

Schenberg, E. E., Alexandre, J. F. M., Filev, R., Cravo, A. M., Sato, J. R., Muthukumaraswamy, S. D.,… & da Silveira, D. X. (2015). Acute biphasic effects of ayahuasca. PloS one, 10(9), e0137202.

Shapiro, K. L., Raymond, J. E., & Arnell, K. M. (1997). The attentional blink. Trends in cognitive sciences, 1(8), 291–296.

Sherwood, A. M., Halberstadt, A. L., Klein, A. K., McCorvy, J. D., Kaylo, K. W., Kargbo, R. B., & Meisenheimer, P. (2020). Synthesis and biological evaluation of tryptamines found in hallucinogenic mushrooms: norbaeocystin, baeocystin, norpsilocin, and aeruginascin. Journal of natural products, 83(2), 461–467.

Spielberger, C. D. (1983). State-trait anxiety inventory for adults.

Stroop, J. R. (1935). Studies of interference in serial verbal reactions. Journal of experimental psychology, 18(6), 643.

Szigeti, B., Kartner, L., Blemings, A., Rosas, F., Feilding, A., Nutt, D. J.,… & Erritzoe, D. (2021). Self-blinding citizen science to explore psychedelic microdosing. Elife, 10, e62878.

Tagliazucchi, E., Zamberlan, F., Cavanna, F., de la Fuente, L. A., Romero, C., Perl, Y. S., & Pallavicini, C. (2021). Baseline power of theta oscillations predicts mystical-type experiences induced by DMT. Frontiers in Psychiatry.

Tellegen, A., & Atkinson, G. (1974). Openness to absorbing and self-altering experiences (“ absorption”), a trait related to hypnotic susceptibility. Journal of abnormal psychology, 83(3), 268.

Timmermann, C., Roseman, L., Schartner, M., Milliere, R., Williams, L. T., Erritzoe, D.,… & Carhart-Harris, R. L. (2019). Neural correlates of the DMT experience assessed with multivariate EEG. Scientific reports, 9(1), 1–13.

Valle, M., Maqueda, A. E., Rabella, M., Rodríguez-Pujadas, A., Antonijoan, R. M., Romero, S.,… & Feilding, A. (2016). Inhibition of alpha oscillations through serotonin-2A receptor activation underlies the visual effects of ayahuasca in humans. European Neuropsychopharmacology, 26(7), 1161–1175.

van Elk, M., Fejer, G., Lempe, P., Prochazckova, L., Kuchar, M., Hajkova, K., & Marschall, J. (2021). Effects of psilocybin microdosing on awe and aesthetic experiences: a preregistered field and lab-based study. Psychopharmacology, 1–16.

Vivot, R. M., Pallavicini, C., Zamberlan, F., Vigo, D., & Tagliazucchi, E. (2020). Meditation increases the entropy of brain oscillatory activity. Neuroscience, 431, 40–51.

Webb, M., Copes, H., & Hendricks, P. S. (2019). Narrative identity, rationality, and microdosing classic psychedelics. International Journal of Drug Policy, 70, 33–39.

Wallach, M. A., & Kogan, N. (1965). Modes of thinking in young children.

Watson, D., Clark, L. A., & Carey, G. (1988). Positive and negative affectivity and their relation to anxiety and depressive disorders. Journal of abnormal psychology, 97(3), 346.

Winstock, A., Barratt, M. J., Maier, L. J., & Ferris, J. (2018). Global drug survey (GDS) 2018. Key findings report.

Yanakieva, S., Polychroni, N., Family, N., Williams, L. T., Luke, D. P., & Terhune, D. B. (2019). The effects of microdose LSD on time perception: a randomised, double-blind, placebo-controlled trial. Psychopharmacology, 236(4), 1159–1170.

